# EFF-1 promotes muscle fusion, paralysis and retargets infection by AFF-1-coated viruses in *C. elegans*

**DOI:** 10.1101/2020.05.17.099622

**Authors:** Anna Meledin, Xiaohui Li, Elena Matveev, Boaz Gildor, Ofer Katzir, Benjamin Podbilewicz

**Affiliations:** Deparitment of Biology, Technion-Israel Institute of Technology, Haifa, 32000, Israel

## Abstract

A hallmark of muscle development is that myoblasts fuse to form myofibers. However, smooth muscles and cardiomyocytes do not generally fuse. In *C. elegans*, the body wall muscles (BWMs), the physiological equivalents of skeletal muscles, are mononuclear. Here, to determine what would be the consequences of fusing BWMs, we express the cell-cell fusogen EFF-1 in these cells. We find that EFF-1 induces paralysis and dumpy phenotypes. To determine whether EFF-1-induced muscle fusion results in these pathologies we injected viruses pseudotyped with AFF-1, a paralog of EFF-1, into the pseudocoelom of *C. elegans*. When these engineered viruses encounter cells expressing EFF-1 or AFF-1 they are able to infect them as revealed by GFP expression from the viral genome. We find that AFF-1 viruses can fuse to EFF-1-expressing muscles revealing multinucleated fibers that cause paralysis and abnormal muscle morphogenesis. Thus, aberrant fusion of otherwise non-syncytial muscle cells may lead to pathological conditions.

**Graphical abstract:** 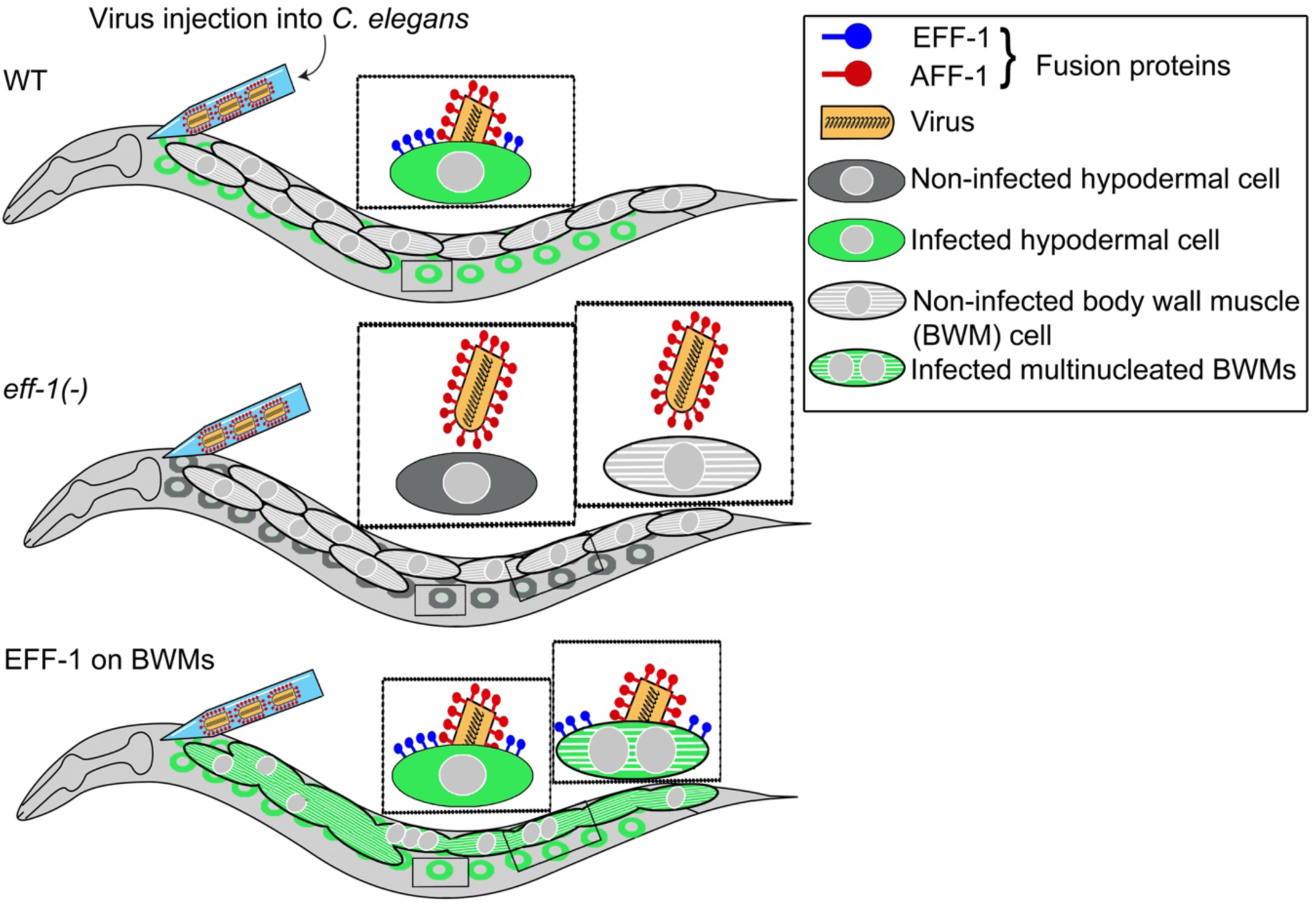

**Significance statement:** Most cells are individual units that do not mix their cytoplasms. However, some cells fuse to become multinucleated in placenta, bones and muscles. In most animals, muscles are formed by myofibers that originate by cell-cell fusion. In contrast, in *C. elegans* the body wall muscles are mononucleated cells that mediate worm-like movement. EFF-1 and AFF-1 fusogens mediate physiological cell fusion in *C. elegans*. By ectopically expressing EFF-1 in body wall muscles we induce their fusion resulting in behavioral and morphological deleterious effects, revealing possible causes of congenital myopathies in humans. Using AFF-1-coated pseudoviruses we infect EFF-1-expressing muscle cells retargeting viral infection into these cells. We suggest that virus retargeting can be utilized to study myogenesis, neuronal regeneration, gamete fusion and screens for new fusogens in different organisms. In addition, our virus retargeting system can be used in gene-therapy, viral-based oncolysis and to study viral-host interactions.

## Introduction

Vesicular Stomatitis Virus (VSV) is an enveloped negative strand RNA rhabdovirus. VSV utilizes its surface glycoprotein G (VSV-G) to infect vertebrates and invertebrates and lyses many cell lines tested to date [1–3]. VSV is widely used for pseudotyping other viruses and has a high transduction efficiency [1–4]. These properties turned VSVΔG into a promising vector for gene therapy, tissue regeneration, viral-based oncolysis, and VSV-based vaccines, some of which successfully completed phase III clinical trials [1,5]. Moreover, VSV-based vectors are useful for studying mechanisms of transcription and replication of RNA viruses, cellular trafficking, antiviral responses and fusion proteins (fusogens) [6–9]. However, the major bottleneck for both applied and fundamental research purposes is achieving an efficient and specific targeting of VSV-based vectors into desired cells/tissues of a live, multicellular organism [3]. Like other viruses, VSV lacks specificity for desired target cells as it suggestively enters cells through highly ubiquitous receptors such as the LDL receptor [10]. Retargeting VSV into particular tissues of interest therefore requires blocking the virus’ natural interactions providing new, cell-specific interactions. Indeed, mutation or substitution of VSV-G with glycoproteins from other viruses or chimeric glycoproteins coupled with antibodies can retarget VSV to specific cells [3,4,11–13].

VSV-G is a class III viral fusogen whereas the *Caenorhabditis elegans* EFF-1 and AFF-1, which fuse somatic cells during development, are structural homologs of class II viral proteins and gamete HAP2(GCS1) fusogens from the fusexin family [14–22]. EFF-1 and AFF-1 also participate in maintaining neuronal architecture and neuronal reconnection following injury [23–27]. Studied viral glycoproteins, including VSV-G, use a unilateral fusion mechanism that depends on the expression of receptors only on the target cells [28,29]. In contrast, EFF-1 and AFF-1 are bilateral fusogens-their presence is required on the membranes of both apposing cells to mediate fusion [6,16,18,30,31]. These two fusogens can act in either a homotypic or a heterotypic manner [6,16,18,21] and mediate heterotypic cell fusion of Sf9 insect and Baby Hamster Kidney (BHK) cells [6,16]. Finally, VSV viruses containing a *GFP* substituting the *VSV-G* coding sequence (VSVΔG) [32,33] that are coated with AFF-1 (VSVΔG-AFF-1) specifically infect AFF-1 or EFF-1 expressing BHK cells [6]. Thus, in contrast to the pseudotyped virus coated with the native, unilateral G glycoprotein (VSVΔG-G), infection by VSVΔG-AFF-1 requires fusogen expression on the host cell membrane. To date, however, VSVΔG coated with AFF-1, EFF-1 or any other non-viral fusogen have not been tested for infection in a living organism. Recently VSVΔG-G was demonstrated to infect living *C. elegans* [7,34]. Given that: (i) VSVΔG-AFF-1 requires a fusogen on the target cell for infection, (ii) a detailed cellular atlas of AFF-1 and EFF-1 expression and function in *C. elegans* is known and (iii) *C. elegans eff-1* and *aff-1* mutants are available, we test whether VSVΔG-AFF-1 can be retargeted to specific cells in living *C. elegans*. We found that AFF-1-coated viruses infect *C. elegans* and specifically target cells that express functional EFF-1 or AFF-1. Furthermore, AFF-1-coated pseudoviruses can be redirected to mononucleated body wall muscles (BWMs) that ectopically express EFF-1. Thus, the new delivery system enabled us to observe that EFF-1-induced BWMs merger and formation of non-functional syncytial muscle fibers, demonstrating the consequences of aberrant muscle fusion in an animal that normally uses mononucleated muscles for gait. Based on our results, we propose that in *C. elegans* a layer of longitudinal mononucleated muscles electrically coupled by gap junctions efficiently mediate wavelike movement and ectopic fusion disrupts the morphology and physiology of the muscular system. We also suggest additional applications for VSVΔG-AFF-1 including finding new fusogens and fusogen-expressing tissues, studying fusogen-fusogen interactions and fusogen-interacting proteins *in vivo*, studying neuronal regeneration processes and using specific cell-delivery approaches for other viruses such as coronaviruses and retroviruses in *C. elegans* and in other organisms.

## Results

### VSVΔG-AFF-1 and VSVΔG-G infect mostly hypodermal and muscle cells respectively

Wildtype, recombinant or pseudotyped VSV has been shown to infect rodents, fish, farm animals, and primates [1,35–37]. Recently, VSVΔG-G was shown to infect *C. elegans* [7]. Previously, pseudotyped VSV expressing the *C. elegans* somatic fusogen AFF-1 (VSVΔG-AFF-1) was shown to infect AFF-1 or EFF-1 expressing BHK cells [6], but was not tested in a living organism. To test whether pseudotyped VSVΔG-AFF-1 [6,33,38] can infect *C. elegans*, we prepared a recombinant VSV strain encoding the fluorescent reporter GFP, coated by AFF-1 fusion protein (VSVΔG-AFF-1). Briefly, BHK cells expressing AFF-1 were infected with VSVΔG-G helper virus (Fig. 1A). Newly generated virions were coated by plasma membrane-bound AFF-1, thereby producing VSVΔG-AFF-1 pseudotyped viruses. Then, we titered the viruses determining the number of viral Infective Units (IU)/ml in BHK-AFF-1 cells. Finally, VSVΔG-AFF-1 viruses were injected into worm’s pseudocoelom; a body cavity filled with fluid that surrounds the internal organs. We expected that virus injected into the pseudocoelom will encounter different cells inside the worm (Figure 1A). Worms were injected with VSVΔG-AFF-1 that was pre-incubated with αVSV-G antibody (see materials and methods and Figure S4), VSVΔG-G (as a positive control) or DMEM medium (as negative control). We used *drh-1(-)* (Dicer Related Helicase -1) mutant worms, as they are more susceptible to VSV infection and do not have any observable phenotypes [7,34]. Taken that VSV-G works unilaterally, while AFF-1 and EFF-1 fusogens have to be present in both fusing membranes [6,16,18,30], we hypothesized that VSVΔG-G and VSVΔG-AFF-1 will produce different infection patterns. Hence, we characterized and compared the GFP positive (GFP(+)) cells infected by VSVΔG-G or VSVΔG-AFF-1. VSVΔG-G preferentially infected muscle cells, including BWM [7], uterine muscles (UM) and stomatointestinal muscles (SM, Figures 1B and 1C). VSVΔG-G also infected the epidermal cells (hypodermis, HYP), excretory canal cell (EC, Figure 1B), glia and neurons in the head (Figures S1A-S1C). In contrast, VSVΔG-AFF-1 infected mostly HYP cells (Figure 1D), and also excretory cell (Figure 1E) pharyngeal muscles (P) (Figure 1D), BWM (Figure 1F) and head neurons and glia (Figures S1D-S1F). VSVΔG-G infects muscles that do not express any known fusogen (67% of infected, GFP(+) worms had only BWM infection), whereas VSVΔG-AFF-1 mainly infects the EFF-1-expressing hypodermis (61% of GFP(+) worms had only hyp infection) (Figure 1G). Moreover, the calculated ID50 (infection dose producing 50% GFP(+) worms) for VSVΔG-G and VSVΔG-AFF-1 were approximately 1000 and 20 IU respectively (Figure 1H). Therefore, VSVΔG-AFF-1, a virus pseudotyped with a bilateral fusogen, can efficiently infect a multicellular organism such as *C. elegans*. Moreover, VSVΔG-AFF-1 and VSVΔG-G are specialized for different cellular targets and have different infectivity levels.

**Figure 1.**
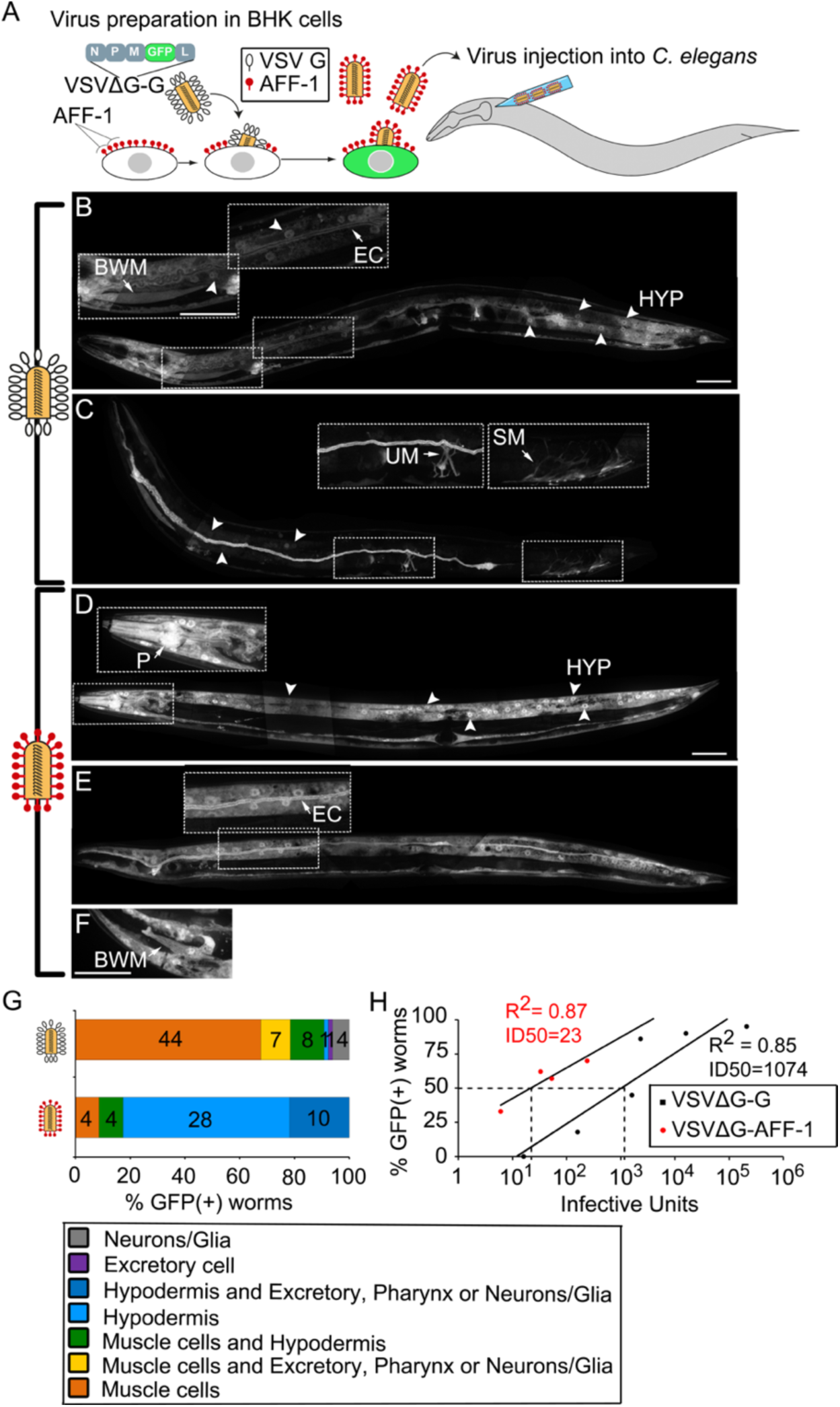
VSVΔG-AFF-1 and VSVΔG-G have different tropisms. **(A)** BHK cells expressing AFF-1 (red pins) were infected with VSVΔG-G. The viral genome encodes GFP replacing VSV-G (white pins). GFP(+) (green), infected cells. VSVΔG-AFF-1 pseudoviruses were harvested from the supernatant, titrated and microinjected into *C. elegans*. **(B-F)** Confocal Z-stack projections of *drh-1* worms injected with VSVΔG-G (2300-16000 IU) or VSVΔG-AFF-1 (33-240 IU) and imaged 48-72 h later (VSVΔG-G, n=65; VSVΔG-AFF-1, n=46 from 5 and 6 independent experiments respectively). Images and their insets are dot boxed. Arrowheads, hypodermal (HYP) nuclei. Arrows, infected cell. BWM-Body Wall Muscle, UM-Uterine Muscle, SM-Stomatointestinal Muscle, EC-Excretory Cell, P-Pharynx. Scale bars, 50 µm. **(G)** Distribution of worms with specified infected, GFP(+) cell types. **(H)** Fraction of GFP(+) worms injected with indicated IU of VSVΔG-G or VSVΔG-AFF-1. Each dot represents an independent experiment (n=10-28 worms per experiment). R^2^ and the IU doses producing 50% GFP(+) worms (ID50) are indicated. See also Figure S1.

### VSVΔG-AFF-1 requires EFF-1 in host cells for infection

We showed that cells infected by VSVΔG-AFF-1 included hypodermis, excretory cell, neurons in the head, glia and pharyngeal cells. It is known that in adults pharyngeal cells, head neurons, glia and hypodermal cells express EFF-1 [14], while pharyngeal cells, excretory duct cell and glia express AFF-1 [18,21] and lastly, excretory cell, head neurons, hypodermal cells and pharyngeal cells express EFF-2, a recent duplication of EFF-1 with unknown function (BG, Oren-Suissa and BP unpublished data). Importantly, BWM cells that do not express any known fusogen in *C. elegans* were infected by VSVΔG-AFF-1 in only 9% of the worms compared to 67% of the worms infected with VSVΔG-G (Figure 1G). These findings suggest that infection of living *C. elegans* by VSVΔG-AFF-1 follows a bilateral action mechanism, and that VSVΔG-AFF-1 is capable of interacting with several members of the fusexin family on the target cell. Therefore, we hypothesized that EFF-1-expressing hypodermal cells could not be infected by VSVΔG-AFF-1 in the absence of EFF-1, while cells expressing AFF-1 or EFF-2 would still be targeted. To test this hypothesis, we injected either VSVΔG-G or VSVΔG-AFF-1 into the pseudocoelom of wt, null *eff-1(ok1021)* mutants *((eff-1(-);*[14]), null *aff-1(tm2214)* mutants ((*aff-1(-)*;[18]) or putative null *eff-2(hy51)* (*eff-2(-)*; see materials and methods) worms, and quantified the percentage of worms with GFP(+) cells. We detected VSVΔG-AFF-1 infected hypodermal cells in *eff-1(+)* (Figure 2A) but not in in *eff-1(-)* worms (Figure 2B), while GFP(+) excretory and glia cell were detected even in the absence of EFF-1 (Figure 2B). *AFF-1(-)* worms had GFP(+) hypodermal cells as in wildtype (Figure 2C). Overall, the fraction of worms showing VSVΔG-AFF-1-infected hypodermal cells was similarly high in wildtype and in *aff-1(-)* (87% and 92% for wt and *aff-1(-)*, respectively) but was absent in *eff-1(-)* mutants (Figure 2D). In addition, muscles were inefficiently infected in wildtype and *aff-1(-)* mutants, and not infected in *eff-1(-)* animals (Figure 2D). The fraction of worms with GFP(+) excretory cells was not significantly different in all backgrounds (Figure 2D). Finally, the fraction of worms with GFP(+) glia/neuron cells was significantly higher in *eff-1(-)*, compared to wt and *aff-1(-)* animals (Figures S1D-S1F). Moreover, *eff-2(-)* and wildtype animals were similarly infected in their hypodermal, muscle and excretory cells (Figure S2). These results show that VSVΔG-AFF-1 requires EFF-1 expressed on hypodermal cells, for their infection. Moreover, we show evidence for AFF-1-EFF-1 bilateral heterotypic interactions *in vivo*.

**Figure 2.**
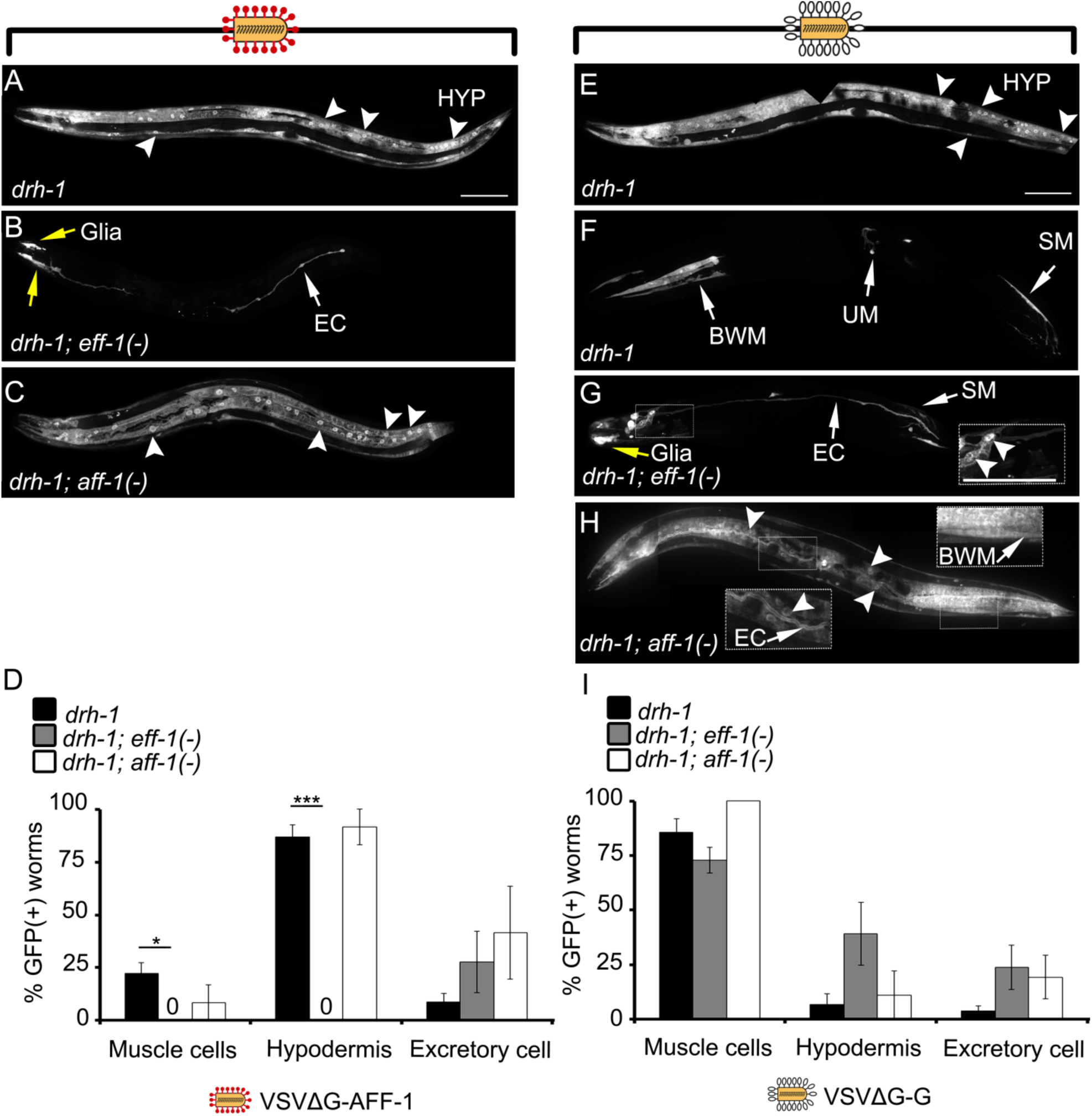
VSVΔG-AFF-1 requires EFF-1 on target cells for infection. **(A-C)** Confocal Z-stack projections of *C. elegans* infected with VSVΔG-AFF-1 (33-240 IU) and imaged 48 h later. *drh-1* (n=46), *drh-1;eff-1(-)* (n=15) and *drh-1;aff-1(-)* (n=10). **(D)** Fraction of GFP(+) worms from (A-C). **(E-H)** Confocal Z-stack projections of *C. elegans* infected with VSVΔG-G (2300-4700 IU) 48 h post-injection. *drh-1* (n=56), *drh-1;eff-1(-)* (n=39) or *drh-1;aff-1(-)* (n=15). **(I)** Fraction of GFP(+) worms from (E-H). Arrowheads, HYP nuclei. Arrows, infected cell. See Figure 1 for cell types. Scale bars, 100 µm. Error bars represent mean ± SEM. *p<0.05, ***p<0.001 (Student’s t-test). See also Figure S2.

EFF-1 mediates formation of hyp7 syncytium by fusing 139 hypodermal cells during embryonic and larval development and *eff-1(-)* worms have mononucleated hypodermal cells instead of this syncytium [14]. To exclude the possibility that hypodermal cells of *eff-1(-)* worms have a reduced susceptibility to viral infection, we injected wt, *aff-1(-)* and *eff-1(-)* worms with VSVΔG-G. We found that wild-type worms injected with VSVΔG-G, presented a full body-length covered by infected hypodermal cells along with muscles and excretory cells (Figures 2E-2F), while *eff-1(-)* worms had only few GFP(+) hyp nuclei localized closer to the injection region-the head, as well as muscles and excretory cells (Figure 2G). *aff-1(-)* worms had hypodermal infection similar to wt animals (Figure 2H). Yet, the fraction of worms injected with VSVΔG-G that showed GFP(+) muscle and hypodermal cells was not significantly different in wt, *eff-1(-)* and *aff-1(-)* backgrounds (Figure 2I). Finally, we observed a small fraction of worms with GFP(+) glia/neurons in the head with no significant difference between all tested backgrounds (Figures S1A-S1C). Thus, hypodermal cells are not compromised for VSVΔG-G infection even in the absence of EFF-1. In addition, VSVΔG-G with its unilateral viral fusogen infects different cells including muscles, hypodermis, excretory cell, and glia/neurons in an EFF-1-and AFF-1-independent manner. This is in contrast to VSVΔG-AFF-1 that uses a bilateral mechanism and infects cells that normally express EFF-1 or AFF-1 in adult animals.

### VSVΔG-AFF-1-mediated infection depends on *eff-1* activity in target cells

We have shown VSVΔG-AFF-1 fails to infect hypodermal cells in *eff-1(-)* animals (Figure 2). To study whether conditional induction of *eff-1* can trigger infection by VSVΔG-AFF-1 we varied the dosage of *eff-1* using temperature shifts in a conditional temperature sensitive (ts) mutant [14,25]. In *eff-1(hy21ts)* animals grown at the permissive 15°C, most epidermal cells fuse to form multinucleated hypodermis and vulva [14]. However, when *eff-1(hy21)* animals are grown at the restrictive 25°C, epidermal cells fail to fuse, producing phenotypes similar to the null *eff-1(ok1021)* worms [16]. Therefore, *eff*-*1(ts)* worms can serve as a temperature inducible system for modification of EFF-1 dosage *in vivo* [14,19,25]. To obtain varying expression of EFF-1, we maintained *eff-1(ts)* animals at 15°C, 25°C or performed 25°C to 15°C downshifts at different developmental stages and then injected VSVΔG-AFF-1 into young adult worms (Figure 3A). Wildtype worms in all experimental conditions had GFP(+) hyp nuclei spanning the worm’s body from head to tail (Figures 3B-3E), while *eff-1(ts)* animals maintained at the restrictive temperature and then shifted down from 25°C to 15°C, had a gradual increase in infected GFP(+) hyp cells (Figures 3F-3I). In wt worms maintained in all conditions, the number of GFP(+) hypodermal nuclei was about 69, consistent with the number of hyp7 nuclei found on one side of the animals body [39,40]. In contrast, for *eff-1(ts)*, the longer time the worms developed at the permissive temperature (15°C), the more GFP(+) hyp nuclei were scored (Figure 3J). Thus, hypodermal infection by VSVΔG-AFF-1 increases with conditional induction of *eff-1*.

**Figure 3.**
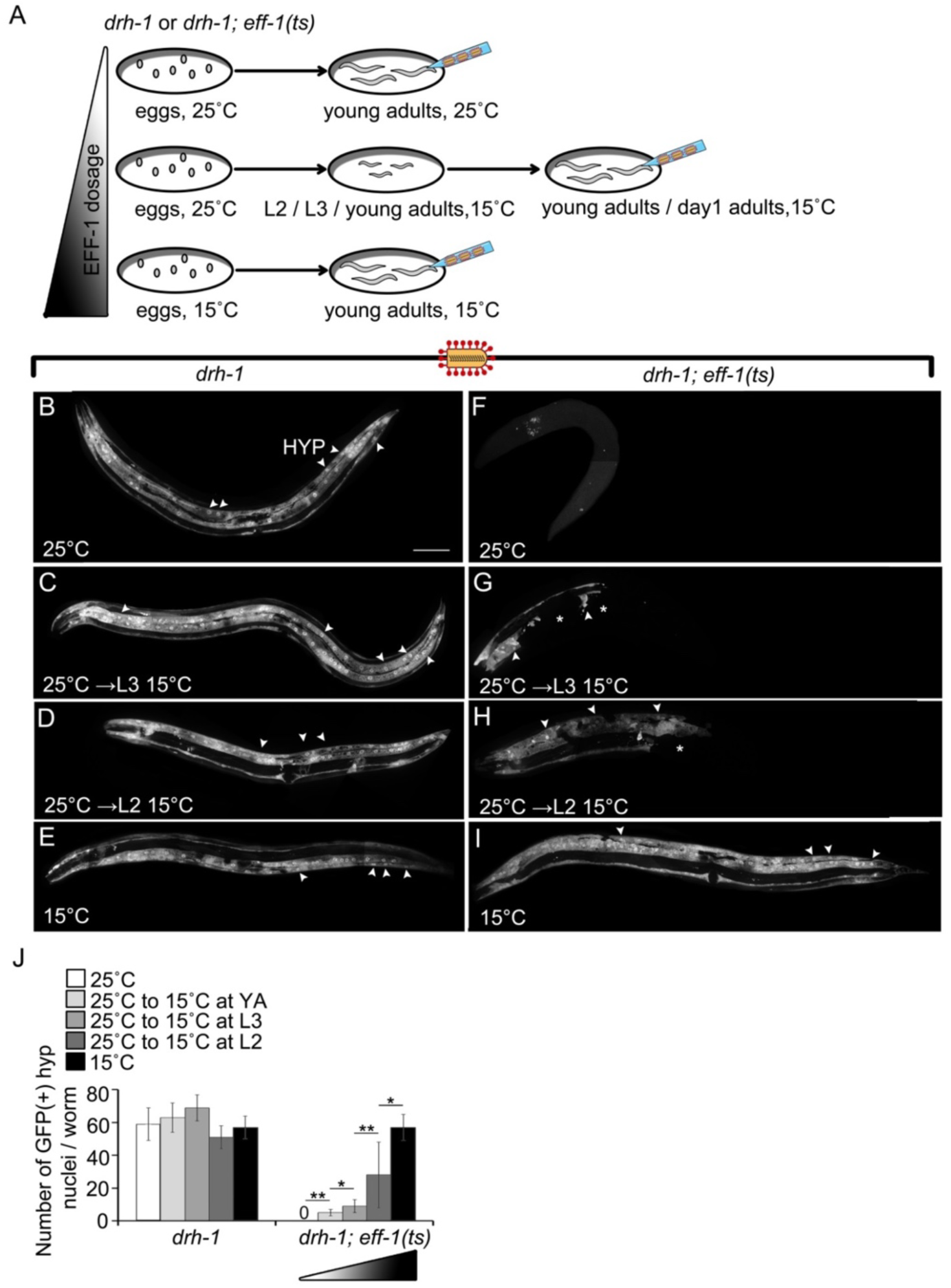
Infection by VSVΔG-AFF-1 increases with induction of EFF-1 function. **(A)** Wt or temperature sensitive *eff-1(ts)* animals were maintained at permissive (15°C) or restrictive (25°C) temperatures for different times during their development. Each row represents a different experimental condition. Blue needles indicate virus injection. L2, Larval stage 2; L3, Larval stage 3. **(B-I)** Confocal Z-stack projections of worms infected with VSVΔG-AFF-1. *drh-1* or *drh-1;eff-1(ts*) animals were injected with VSVΔG-AFF-1 (7-125 IU) as adults, and imaged 48 h later. Arrowheads, HYP nuclei. Asterisks, patches of GFP(-) surrounded by GFP(+) hypodermal cells. Scale bar, 100 µm. **(J)** Quantitation of experiments. Error bars average ± SEM. Student t-test. *p<0.05, **p<0.001. n=3-10 worms per condition. Triangle, dosage of *eff-1*.

### Ectopic EFF-1 expression in BWMs produces dumpy and uncoordinated worms

We demonstrated that VSVΔG-AFF-1 specifically infects EFF-1 expressing cells and rarely infects muscle cells. In addition, EFF-1-dependent infection could be induced by EFF-1(ts) conditional expression in target cells (Figure 3). Hence, we propose that ectopic EFF-1 expression in host cells, such as body wall muscles (BWMs), could retarget VSVΔG-AFF-1 into these cells. *C. elegans* has 95 mononucleated rhomboid BWMs, arranged as staggered pairs and bundled in four quadrants that run along the worm’s body [41–43]. While most vertebrates and invertebrates have syncytial striated muscles composed of long multinucleated myofibers, in *C. elegans* and other nematodes the BWMs have not been described to fuse and do not form multinucleated myofibers (https://www.wormatlas.org/hermaphrodite/muscleintro/MusIntroframeset.html). Therefore, we hypothesized that EFF-1 expression in BWMs can induce their ectopic fusion, alter their structure, produce muscle-related phenotypes and increase VSVΔG-AFF-1 infection of BWMs. First, we monitored animals with an extra-chromosomal array containing genomic EFF-1 under the muscle-specific *myosin-3* promoter (*myo-3p*::EFF-1), together with *myo-3p::*mCherry which labels cytoplasm and nuclei in BWMs, enteric, gonadal and vulval muscles. Most adult animals, including *myo-3p*::mCherry (+) and (-), were wild-type-like (Table S1, Figures 4A-4B and Movie S1), while about 10% of animals were both *myo-3p*::mCherry(+) and had an uncoordinated and dumpy (Unc+Dpy) phenotype (Table S1, Figures 4C-4D and Movie S1). Most *myo-3p*::mCherry(+), Unc+Dpy animals arrested during larval development (Table S1) with additional phenotypes including bridged BWMs behind the terminal bulb (0/12 wt-like worms and 9/15 Unc+Dpy worms), mCherry-labeled aggregates that could be observed with DIC (0/12 wt-like worms and 9/15 Unc+Dpy worms) (Figures 4E-4H), *myo-3p*::mcCherry(+) BWMs with clustered nuclei (3/12 wt-like worms and 9/15 Unc+Dpy worms) and elevated number of nuclei/ BWM cell (1.4±0.2 and 2.2±0.3 nuclei/BWM cell in wt-like vs Unc+Dpy larvae respectively) (Figures 4E-4I). While ectopic *hsp*::EFF-1 expression causes embryonic lethality [14,19,25], we found no difference between the fractions of *myo-3p*::mCherry (+) and (-) unhatched eggs (Table S1). Thus, EFF-1 expression in BWMs accounts for the Unc+Dpy phenotypes, larval arrest and clustered BWM nuclei, which supports the hypothesis that ectopic EFF-1 induces BWMs multinucleation by cell-cell fusion causing behavioral and morphological phenotypes.

**Figure 4.**
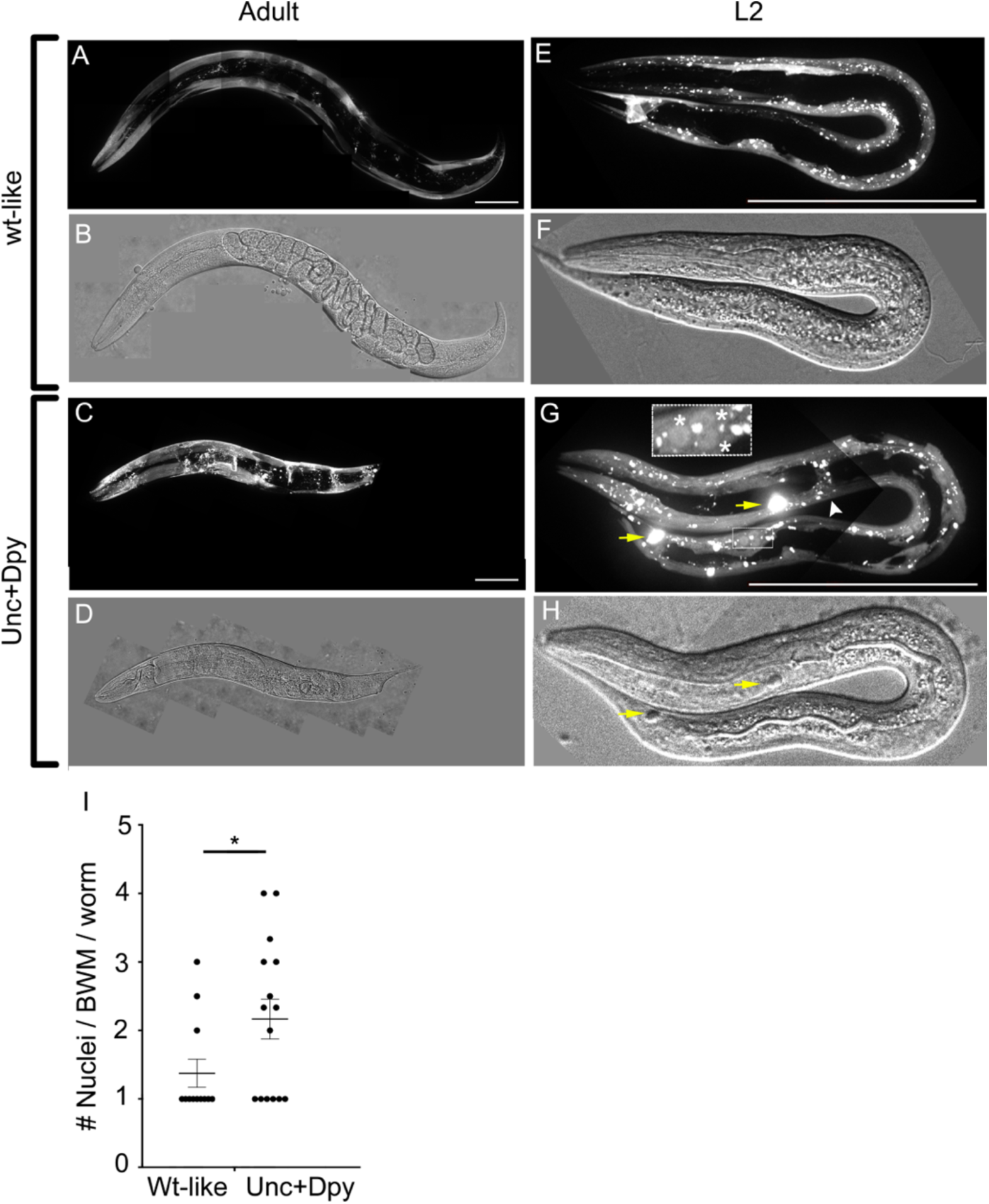
EFF-1 expression in BWMs produces Unc+Dpy worms with multinucleated cells. **(A-H)** Images of fluorescent Z-stack projections and respective DIC of animals with extrachromosomal *myo-3p*::EFF-1, *myo-3p*::mCherry. (G) White arrowhead, bridge formed between 2 BWMs from opposing quadrants. Yellow arrows, *myo-3p*::mCherry accumulations also in DIC (H). Asterisks, clustered nuclei within one BWM. Scale bars, 100 µm. **(I)** Number of nuclei per *myo-3p*::mCherry (+) BWM cell in L2s. wt-like (n=12); Unc+Dpy (n=15). Each dot represents the average number of nuclei/BWM cell/worm. Total average ± SEM for each phenotype. Two tailed Student’s t-test p<0.05*. See also Table S1.

### EFF-1 expression in BWMs induces their fusion and retargets VSVΔG-AFF-1 to muscles

To determine whether EFF-1 expression in BWMs can induce their fusion, we imaged BWMs expressing a membrane-bound YFP (MB::YFP), *myo-3p*::EFF-1 and *myo-3p*::mCherry, (see materials and methods). BWMs of wt-like worms have a normal spindle-shaped morphology with cellular membranes between adjacent BWMs showing different levels of cytoplasmic *myo-3p*::mCherry (as *myo-3p*::mCherry is an extrachromosomal array) (Figures 5A-5C). In contrast, Unc+Dpy worms had disordered muscle fiber structure, the membranes surrounding the BWMs were indistinguishable and the BWMs showed an evenly distributed cytoplasmic *myo-3p*::mCherry indicating fusion and content mixing between these cells during development (Figures 5D-5F). Thus, EFF-1 expression in BWMs induces disappearance of cellular membranes and cytoplasmic merger, suggesting that EFF-1-mediated cell-cell fusion alters BWM structure, likely resulting in Unc+Dpy and larval arrest phenotypes.

**Figure 5.**
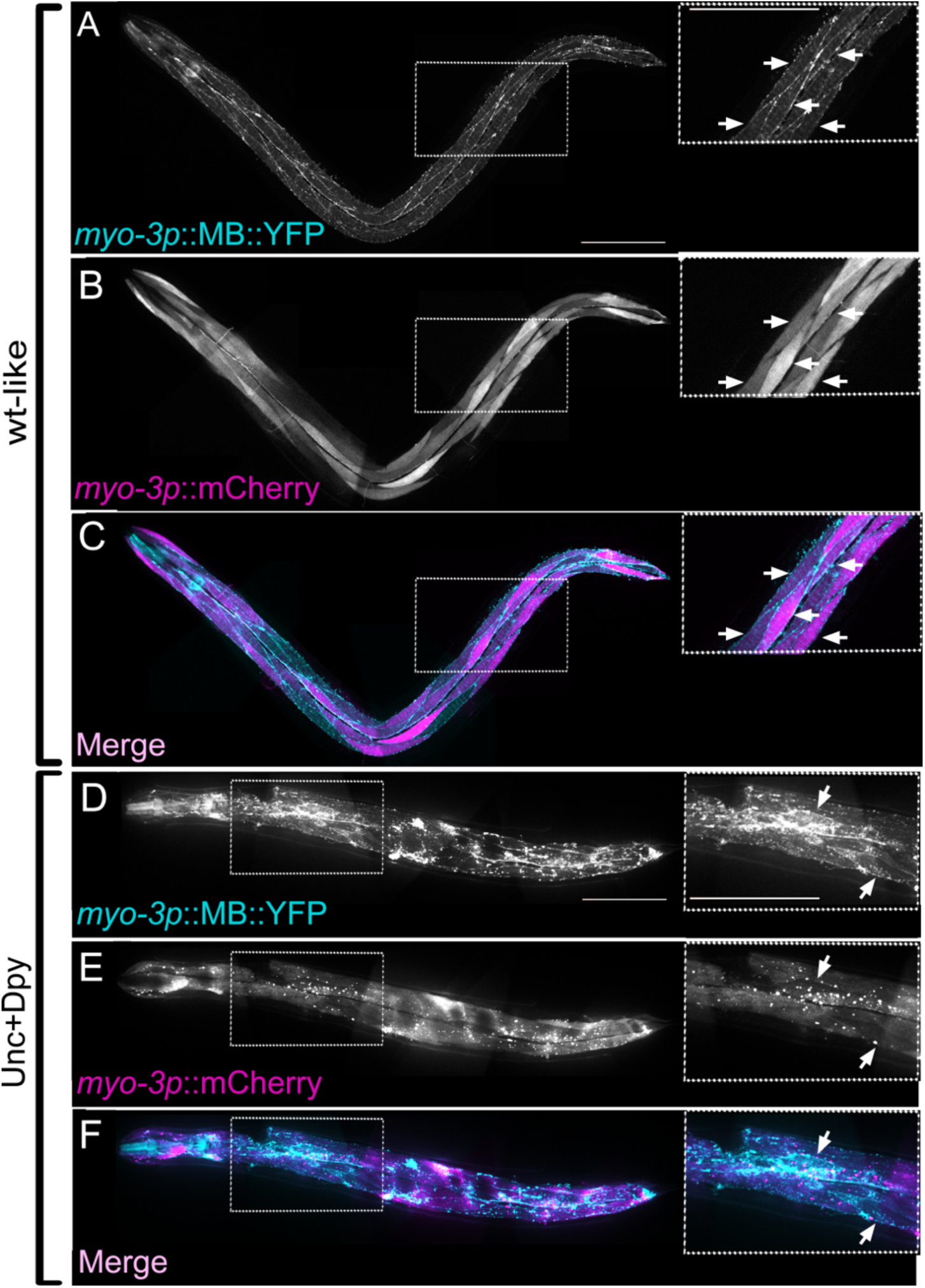
EFF-1 expression in BWMs induces their fusion. (**A-C**) Confocal images of wt-like adult worms with membrane bound (*MB*) *myo-3p::MB::YFP* (cyan) and extrachromosomal array containing *myo-3p::*EFF-1, *myo-3p::*mCherry (magenta). (**D-F**) Confocal images of Unc+Dpy [*myo-3p::MB::YFP* (cyan); *myo-3p::EFF-1, myo-3p::mCherry]*. Arrows, unfused BWMs with MB (cyan). Note only two unfused BWMs, all the others appear fused with no MB separating them. Insets correspond to white-dotted area. Scale bars 100 µm. See also Movie S1.

We showed that VSVΔG-AFF-1 infects EFF-1-expressing hypodermal cells in >80% of the infected worms but infects muscle cells that do not express EFF-1 in about 20% of the infected animals (Figure 2D). Moreover, EFF-1 expression in BWMs induced their fusion, and generated Unc+Dpy worms (Table S1, Movie S1 and Figures 4A-4D and 5A-5F). Taken that EFF-1 and AFF-1 are bilateral fusogens and VSVG is a unilateral fusogen, we hypothesized that ectopic expression of EFF-1 in BWMs would increase their infection by VSVΔG-AFF-1 but not by VSVΔG-G. Therefore, we injected either VSVΔG-AFF-1 or VSVΔG-G into the pseudocoelom of worms ectopically expressing EFF-1 and mCherry in BWMs and imaged them. For VSVΔG-AFF-1 but not for VSVΔG-G, the fraction of infected BWMs was significantly higher in Unc+Dpy worms (expressing the *myo-3p*::mCherry marker) compared to wt-like (expressing the *myo-3p*::mCherry marker) and wt animals (Figure 6). Thus, EFF-1 expression in BWMs retargets VSVΔG-AFF-1 into muscles but does not affect the natural VSVΔG-G tropism for BWMs.

**Figure 6.**
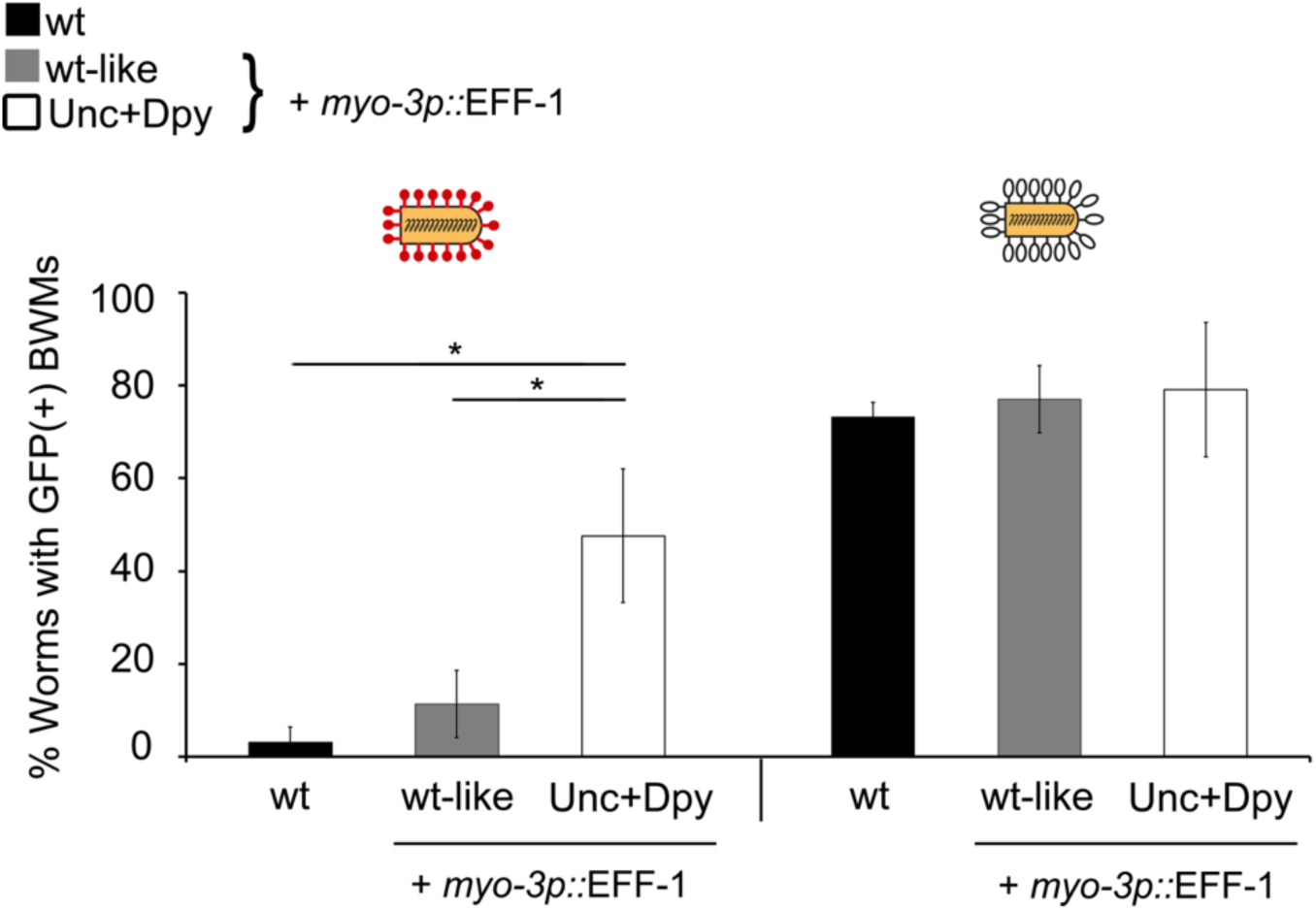
Retargeting of VSVΔG-AFF-1 to body wall muscle cells. Wild-type worms and animals with extrachromosomal array containing *myo-3p::*EFF-1 and *myo-3p::*mCherry were injected with VSVΔG-AFF-1 (35-63 IU, red pins; n=39 wt, n=50 wt-like and n=27 Unc+Dpy worms) or VSVΔG-G (3*10^5^ IU, white pins; n=30 wt-like and 14 Unc+Dpy) respectively. Wt worms injected with VSVΔG-G (2300-4700 IU, n=56) were taken from figure 2I. Animals were analysed by SDC microscopy. Data represents average percentage of worms with GFP(+) BWMs ± SEM. Student’s t-test: *p<0.05.

We showed that VSVΔG-AFF-1 infects neuron and glia cells in the head (Figures S1D-S1F). In order to assess the infection of other fusogen-expressing neurons, we chose the arborized neuron pair PVD, previously shown to express EFF-1, which spans the length of the worm [25], VSVΔG-AFF-1-infected PVD cells were not observed. Nevertheless, we found 3 wt animals (out of >100 infected worms), where PVD cells were infected with VSVΔG-G following injections with high dosage of pseudotyped viruses (Movie S2). Moreover, we tested whether ectopic AFF-1 expression in PVD or ectopic EFF-1 expression in 12 mechanosensory and chemosensory neurons in *eff-1(ts)* background at the restrictive 25°C, could induce VSVΔG-AFF-1 infection in these cells. None of these treatments produced VSVΔG-AFF-1 infection in the neurons ectopically expressing EFF-1 or AFF-1 (Figure S3 and Table S2). There are several possibilities to explain why VSVΔG-AFF-1 did not infect these neurons; (i) relatively low titer of VSVΔG-AFF-1 preps, (ii) a physical barrier such as hypodermis, sheath cells or glia cells that surround neurons or (iii) diffusion of the GFP signal in long neuronal processes.

### EFF-1 expression in fused BWMs enables VSVΔG-AFF-1 and VSVΔG-G spreading

Based on the aberrant muscle fusion phenotype seen in Unc+Dpy worms following EFF-1 expression in BWMs, we hypothesized that viral spreading through such novel BWM syncytia would be enhanced. To test this hypothesis, we injected VSVΔG-AFF-1 or VSVΔG-G into wt-like or Unc+Dpy worms and quantified the number of BWM cells that were either (i) mCherry(+) only (expressing EFF-1 and *myo-3p*::mCherry(+)), (ii) GFP(+) only (infected but not expressing EFF-1) or (iii) mCherry(+) and GFP(+) expression in the same cells (infected and expressing EFF-1). We found that for both VSVΔG-AFF-1 and VSVΔG-G, wt-like worms had individual GFP(+) BWM cells, that were mostly mCherry (-) (Figures 7A-7F, arrows), while Unc+Dpy animals had continuous GFP(+) BWMs that overlapped with mCherry along their body length (Figures 7G-7L, dashed lines and arrows). Compared to wt-like animals, the Unc+Dpy worms had about five-times fewer non-infected mCherry(+) only BWMs, a similar number of infected-GFP(+) only cells and about fifteen-fold more infected BWM cells that also express mCherry (Figures 7M-7N). Lastly, Unc+Dpy worms had ∼2.5 nuclei per VSVΔG-AFF-1-or VSVΔG-G-infected BWMs, compared to wt-like animals with 1 nucleus per BWM cell (Figures 7O-7P). Therefore, EFF-1 expression in BWMs transforms mononucleated BWMs into syncytial muscle fibres enabling viral spreading within the fused cells and increasing the number of both VSVΔG-AFF-1-and VSVΔG-G-infected BWM cells.

**Figure 7.**
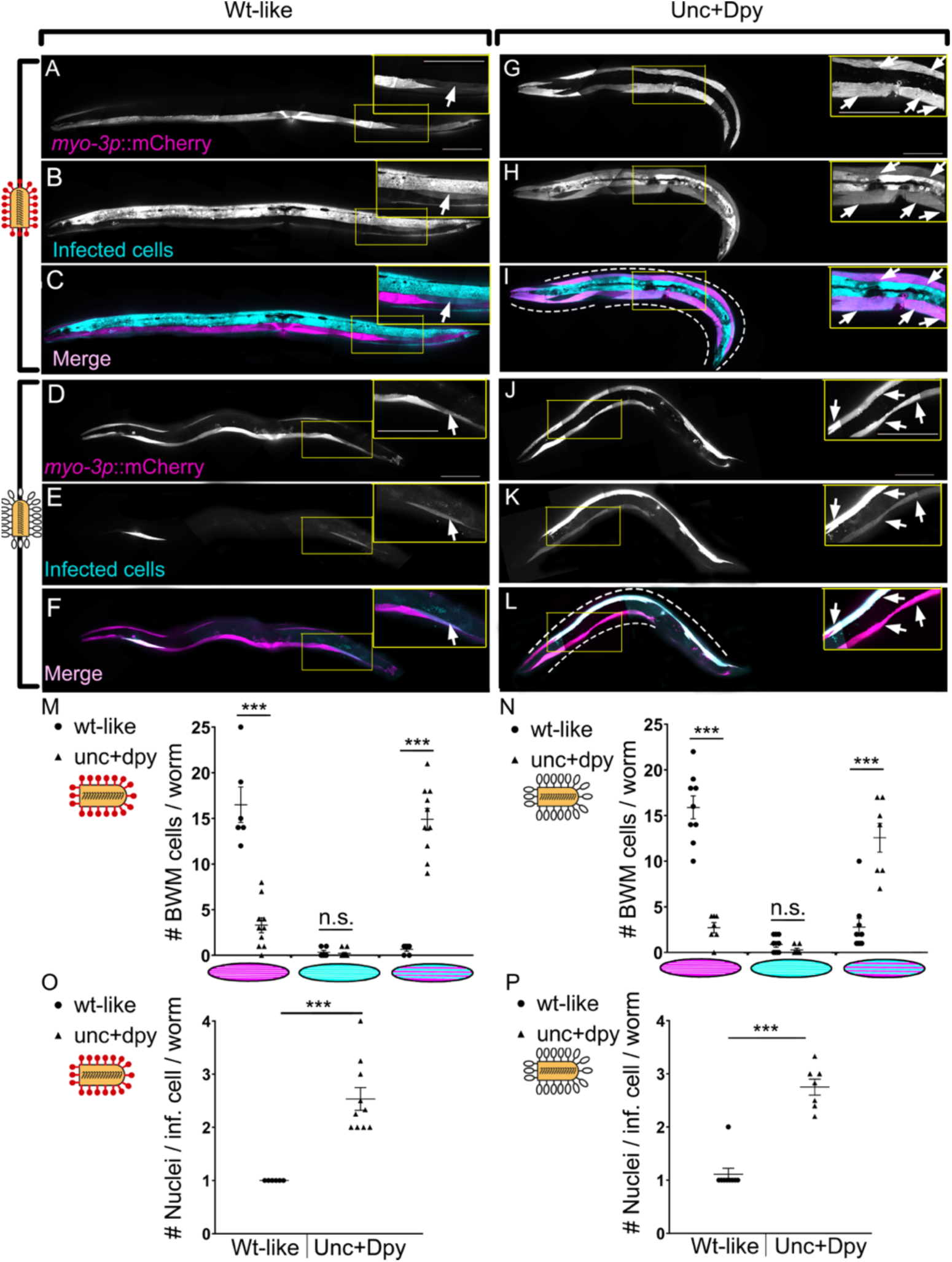
EFF-1 expression in BWM enables VSVΔG-AFF-1 and VSVΔG-G spreading along fused muscles. **(A-L)** Z-stack projections of wt-like (A-F) and Unc+Dpy animals (G-L) expressing *myo-3p::*EFF-1 and *myo-3p::*mCherry infected with VSVΔG-AFF-1 (35-63 IU red pins) or VSVΔG-G (3*10^5^ IU; white pins). Insets and their corresponding images (yellow frames). Arrows, individual infected (cyan) BWMs. Dashed lines outline grouped BWMs that express *myo-3p*::mCherry and *myo-3p::*EFF-1 (magenta) and infected with virus (cyan) showing spreading of GFP. Scale bars, 100 µm. **(M-N)** Number of BWM cells/worm expressing EFF-1 (magenta cell), infected (cyan cell) or expressing EFF-1 and infected (magenta and cyan). wt-like (circles) and Unc+Dpy (triangles). Each point represents a single worm. **(O-P)** Quantitation of multinucleation of infected BWMs. Each dot represents an average number of nuclei/ GFP(+) BWM, calculated from 1-6 multinucleated BWMs of a single worm. (M and O) wt-like n=6 and Unc+Dpy n=10 animals. (N and P) wt-like n=9 and Unc+Dpy n=7 animals. Black horizontal lines, average ± SEM. Student’s t-test, *** p<0.0001; n.s., not-significant.

## Discussion

### Utilizing VSVΔG-AFF-1 to study muscle fusion and muscle fusogens

Myogenesis of striated muscle involves the formation of multinucleated myofibers by myoblast-myoblast fusion. Myoblast fusion is also essential for muscle tissue patterning, maintenance and damage repair [62,63]. In contrast, *C. elegans* BWMs do not fuse and exist as mononucleated cells, contracting in synchrony that mediate gait [64]. We used VSVΔG-AFF-1 to label EFF-1-expressing BWMs and found that EFF-1-mediated BWMs fusion results in loss of coordination, although we did not determine whether the observed locomotory defects result from aberrant joining of separately-innervated muscles or from a loss of contractility. These results demonstrate that a locomotory circuit may be optimally adjusted for both separate (as in *C. elegans*) and syncytial (as in vertebrate) striated muscles. Indeed, failure of myoblast fusion in vertebrates may have equally severe phenotypes [47,48]. Despite the importance of muscle fusion, the studies of muscle fusion proteins are just emerging [55–58]. We suggest that VSVΔG-AFF-1 can be used in screens to identify additional muscle fusogens in invertebrates (e.g. Drosophila). Alternatively, viruses coated with the bilateral Myomaker, can be a tool to screen for muscle-expressed Myomaker-interacting proteins in vertebrates.

### VSVΔG-AFF-1 infects, and can be retargeted to muscles expressing EFF-1 or AFF-1

Previously described viral-vector-based systems used viral unilateral fusogens to target desired cells. For instance, fusogens that are either native, modified or from different viral origin, change the specificity of the viruses targets and facilitate the delivery process [1–3]. In addition, a potent targeting of cancer cell lines was achieved with adenovirus (non-enveloped virus) armed with an adaptor composed from 2 designed ankyrin repeat proteins (DARPins), one binding to the viral fiber knob and the second binding to cancer-specific markers [44]. Our work presents a new approach to retarget viral vectors into muscle cells, based on viruses coated with a nematode bilateral fusogen, namely AFF-1 and expression of bilateral fusogens such as EFF-1/AFF-1 in desired host cells. VSVΔG-AFF-1 has a high degree of specificity towards cells that express EFF-1/AFF-1 and can be redirected by expressing them on a certain cell (e.g. BWMs). Moreover, VSVΔG-AFF-1-mediated infection can be manipulated quantitatively by changing the amount of a partner fusogen expressed on the target cells. VSV and VSV-pseudotypes successfully infect invertebrate and vertebrate cell-lines and model organisms [1,2,35–37]. Although EFF-1 and AFF-1 are not present, and have no known homologs in vertebrates, VSVΔG-AFF-1 infects mammalian BHK cells expressing AFF-1/EFF-1 [6]. It will be interesting to test if VSVΔG-AFF-1 can specifically target cells ectopically expressing AFF-1/EFF-1 in a vertebrate model. Similarly, pseudotyped VSVΔG-SARS-CoV-2-S-glycoprotein [4] could be targeted to humanized worms or mice expressing hACE2 to study entry mechanisms, pathogenesis and potential treatments for viral infections, including COVID19 in model organisms.

### Additional applications for VSVΔG-AFF-1

In the recent decade, extracellular vesicles (EVs) emerged as intercellular communication carriers, containing variable cargoes (e.g lipids, DNA, RNA, toxins and proteins) and involved in different biological processes including cell-cell interactions, cancer, tissue and neuronal regeneration. EVs can serve as biomarkers and are potent vehicles for gene and drug delivery [26,45,46]. However, work with EVs-based vectors has to tackle the issues of specific targeting, loading of desired cargo, presence of undesired/non-specific content and quantification of produced EVs. Our system consisting of enveloped pseudotyped virus coated with bilateral fusogens and host muscle cells expressing bilateral fusogens, demonstrates cell-specific targeting. Moreover, VSVΔG-AFF-1 has the benefits of viral-based vectors, namely loading a specific and tailored cargo including fluorescent markers such as GFP-coding RNA sequence, simple vector quantification and cargo amplification within the host cell. Therefore, VSVΔG-AFF-1 may serve as a paradigm for EVs-mediated transport. It can be utilized to explore how EVs deliver their cargos and to study whether and how EVs are involved in different biological processes.

Exoplasmic membrane fusion is a fundamental biological process involved in myogenesis, fertilization, vulva development, bone formation and resorption, hypodermis morphogenesis, placentation, tubulogenesis, neuronal regeneration and viral infection [28,31,47,48]. Despite the importance of these processes, very few cell-cell fusogens have been identified and characterized so far. These include: (i) Syncytins involved in placenta formation [49–51]; (ii) Fusexins including EFF-1 and AFF-1 from *C. elegans* [14,17–22,31] and HAP2/GCS1 which mediates gamete fusion in protists and flowering plants [38,52–54] and (iii) Myomaker (TMEM8c) and myomerger (myomixer/minion) proteins that fuse muscle cells in vertebrates [55–58]. We demonstrate that VSVΔG-AFF-1 injected into *C. elegans* encounters and infects different EFF-1/AFF-1-expressing cells. Ectopic expression of EFF-1 in BWMs induces these mononucleated muscles to form syncytial myofibers, however, these multinucleated muscles are pathological and the worms become deformed and paralyzed. Moreover, bilateral heterotypic fusion has been described between EFF-1 and AFF-1 and also between EFF-1 and sperm HAP2/GCS1 from *Arabidopsis* expressed on two populations of BHKs [38]. Therefore, VSVΔG-AFF-1/EFF-1 may be exploited in screens to identify new bilateral fusogens or fusogen-interacting proteins in different organisms and to understand

how and why some myoblasts fuse (e.g. skeletal muscles) while others function optimally as individual mononucleated cells (e.g. smooth muscles and BWMs).

## Materials and methods

### Nematode strains

Unless otherwise stated, all nematodes were maintained at 20°C, according to standard protocols [65,66]. *drh-1(tm1329)* served as the wild type background. Mutations and strains that were used in this study are listed in Table S3.

### DNA constructs

The *myo-3p*::EFF-1 plasmid was constructed by cloning the *myo-3* promoter region from *myo-3p*::mCherry plasmid with Sal I (New England BioLabs Cat#R3138) and Nhe I (ThermoFisher Cat# FD0974) and inserting it into the *hsp16-2*:: EFF-1 plasmid cut with the same enzymes to replace the original heat shock promoter. To produce *eff-2 (hy51)* mutant worms with CRISPR, the following DNA constructs were generated:

1. pBG115 plasmid encoding single guide targeting *eff-2* to insert *mNeonGreen* was generated by cloning *eff-2* targeting sequence into the CAS9 plasmid pDD162 with BG84 and BG85 primers.
2. BG123 conversion oligonucleotide encoding wt *pha-1* fragment was PAGE purified.
3. PCR amplicon encoding for the mNeonGreen and unc-54 3’UTR flanked by homology arms to the *eff-2* gene, generated by amplification of mNeonGreen from plasmid X, with BG129 and BG130 primers containing a 50 bp homology arms to *eff-2*.

Sequences of primers and oligonucleotides mentioned above are found in Table S3.

### Transgenic animals

For standard extrachromosomal transgenes, germline transformation was performed using standard protocols [67]. Transgenic lines were kept as extrachromosomal arrays and maintained by following the expression from either *myo-3p*::mCherry, *myo-2p*::GFP, or *mec-4p*::dsRed, *odr-1p*::dsRed plasmids that were co-injected as transformation markers. *myo-3p::*mCherry encodes mCherry expression in body wall muscle and vulva muscle cells. *myo-2p*::GFP encodes GFP expression specifically expressed in pharyngeal muscles. *pmec-4*::dsRed encodes dsRed expression in six touch receptor neurons. *odr-1p*::dsRed encodes dsRed expression in two odor sensory neurons. Plasmids mentioned above and strains containing these arrays are listed in Table S3. Transgenic lines with extrachromosomal arrays were generated as follows:

- BP2126: *drh-1(tm1329);eff-1(hy21)* injected with 20ng/µl *pmyo-2::GFP* as transformation marker and 0.1ng/µl *pdes-2::AFF-1*.
- BP2131-3: *drh-1(tm1329);eff-1(hy21)* injected 10ng/µl *pmec-4::dsRed* and 10ng/µl *podr-1::dsRed* as transformation markers and to label sensory neurons, 1ng/µl *pmec-4::EFF-1* and 1ng/µl *podr-1::EFF-1*.
- BP2137: *drh-1(tm1329)* injected 10ng/µl *myo-3p::mCherry* as transformation marker and to label Body Wall Muscle cells, and 1ng/µl *myo-3p::EFF-1*.

*eff-2(hy51)* allele was generated by CRISPR/Cas9 [65,66] insertion of mNeonGreen into the first exon of *eff-2* gene. *pha-1* was used as a conversion marker. *pha-1(e2123)* temperature sensitive worms were injected with 50 ng/µl pBG115 targeting *eff-2* and containing CAS9, 50 ng/µl pJW1285 sgRNA plasmid against *pha-1* that contains the CAS9, 20 ng/µl BG123 conversion oligonucleotide encoding wt *pha-1* fragment and 20 ng/µl amplicon encoding for the mNeonGreen and *unc-54* 3’UTR flanked by homology arms to the *eff-2* gene. Worms that survived development at the restrictive temperature (25°C) were screened and sequenced for mutations in the *eff-2* gene. *hy51* is the result of an imprecise partial inverted insertion of the PCR fragment into the *eff-2* locus. One bp was deleted at position +31 and 488bp of the unc-54 3’UTR and mNeonGreen gene were inserted into *eff-2* coding region. The product of the inverted insertion is a non-functional fluorescent protein that causes a frame shift in *eff-2* and terminates its translation at Amino Acid #18. The *eff-2* targeting sequence was added to primers (see Table S3) and inserted to the linearized vector using restriction-free cloning technique.

### Live imaging of worms

For imaging of viral infection and BWM fusion in *C. elegans*, worms were analyzed by Nomarski optics and fluorescence microscopy using Nikon eclipse Ti inverted microscope with Yokogawa CSU-X1 spinning disk confocal (SDC) as described previously [26,27]. Briefly, animals were anesthetized in 0.01-0.05% tetramisole in M9 solution for 20-30 min and then picked with an eyelash attached to a toothpick and transferred to a 5 µl droplet of M9 solution placed on 3% agar slide. Images were acquired with Metamorph software, when using the spinning disk confocal. Z-stacks were taken with Plan Fluor 40x NA=1.3 or Apochromat 60x NA=1.4 objectives. Excitation of GFP was achieved with 488 nm wavelength laser (2-8% intensity, 100 ms exposure time). mCherry was excited with 561 nm wavelength (15-20% intensity, 100 ms exposure time). ∼0.5 μm z-steps were recorded with iXon3 EMCCD camera (Andor). Multidimensional data were reconstructed as maximum intensity projections using Fiji software (NIH ImageJ). For live imaging of worms with fused BWM phenotypes, plates with worms were placed on the stage of Zeiss stereo Discovery V8 stereo microscope. Images and movies were captured at x8 magnification with additional magnification from PlanApo S 2.3X objective and a Hamamatsu ORCA-ER camera controlled by micromanager software (https://micro-manager.org). Figures were prepared using Fiji, Adobe Photoshop CS5 and GraphPad Prism 8.

### DNA transformation and viral infection by microinjection

Microinjections were performed as described [7,34] with some modifications. Shortly, late L4 or young adult worms were placed into droplet of halocarbon oil on 3% agarose pads. Pulled capillary needles were secured onto a Nikon DIAPHOT 300 microscope equipped with a micromanipulator and regulated pressure source (Narishige). For DNA transformation, needles were loaded with 0.8 µl with DNA solution containing TE buffer, the target construct DNA, a co-injection marker DNA and the required amount of with pKSI-1 (empty vector) DNA to reach a total concentration of 100 ng/µl DNA. For experiments with viral infection, needles were loaded 0.8 µl DMEM (mock-infections) or pseudo viruses in DMEM+5%FBS. Infection doses of VSV are based on VSV titration on BHK cells and used 10nL volume as a single microinjection dose [7]. Agar pads with worms were placed on stage of Nikon DIAPHOT 300 microscope. Worms were observed under x40 objective. For DNA transformation, animals were injected into the gonad and immediately placed into droplet of M9 buffer placed on NGM plates seeded with OP50-1 *E. coli*. For experiments with viral infection worms were injected into pseudocoelom -behind the terminal bulb of the pharynx and were immediately placed into droplet of M9 buffer placed on NGM plates with 50 μg/mL FUdR (Sigma), seeded with OP50-1 *E. coli*. Unless otherwise stated, animals were maintained at 25°C until scoring of infection.

### Scoring viral infection

Scoring viral infection using fluorescence assay was performed as described [7,34] with some modifications. Briefly, 48-72 hours post injection worms were processed for live imaging as described above. Animals that were unresponsive to prodding by a platinum wire worm pick were considered dead and were removed from the experiment. Animals that crawled off the plate or were lost during the experiment were censored. Worms with ≥1 GFP (+) cells were considered as infected worms, while not injected or DMEM injected worms served as a negative controls. For different experiments, either the number of GFP(+) cells/animals or the type of infected tissues were observed and quantified.

### Temperature shift experiments

Temperature sensitive *eff-1(hy21ts)* mutant worms were synchronized by hypochlorite treatment of adult worms. The obtained eggs were left for overnight L1 hatching on NGM plates without food at 20° C. L1 animals were then transferred to NGM plates with OP50 bacteria at 15° C or 25° C incubators until reaching the desired developmental stage. The plates were either downshifted from 25° C to 15° C, or left at 25° C or at 15° C throughout the experiment. The developmental stages were determined by analyzing gonadal size and structure using Nomarski optics. Finally, worms were injected with VSVΔG-AFF-1, maintained and imaged as described above.

### Counting overlapping cells

To test if EFF-1 expressing BWMs were infected by VSVΔG-AFF-1, we utilized worms with an extrachromosomal array expressing EFF-1 under the *myo-3* promoter and a plasmid containing *myo-3p*::mCherry. We found that most mCherry (+) worms had a wt-like phenotype, but a small fraction of these worms were Unc+Dpy. Importantly, all sibling mCherry(-) worms were wt-like (Table S1). Hence, for the experiment we utilized a fluorescence stereomicroscope and chose late L4/ young adult, wt-like and Unc+Dpy animals that had continuous *myo-3p::mCherry*(+) BWMs (which could express EFF-1 and fuse with each other). Next, these worms were injected with VSVΔG-AFF-1 or VSVΔG-G. 48-72h later, Z-stack images of the worms were obtained with SDC microscope (as described in live imaging section). Worms with ≥1 infected BWM were selected. For each worm, we counted the number of BWMs that were mCherry(+) only (express EFF-1), GFP(+) only (infected by virus) and overlapping-with both mCherry(+) and GFP(+) (express EFF-1 and infected).

### Counting number of nuclei per BWM

To test if ectopic EFF-1 expression in BWMs produces multinucleated BWMs, wt-like and Unc+Dpy animals that had continuous *myo-3p::mCherry*(+) BWMs (which could express EFF-1 and fuse with each other) were imaged by SDC microscope as described above. In one set of experiments we imaged L2 worms and counted number of nuclei per *myo-3p::mCherry* cell. In a second set of experiments, we imaged adult worms with BWMs infected by either VSVΔG-AFF-1 or VSVΔG-G and counted number of nuclei per infected (GFP(+)) BWM cell. For both sets of experiments, in worms with at least one multinucleated BWM we monitored all multinucleated BWMs and calculated the average number of nuclei/BWM for each worm. Worms in which multinucleated cells were not observed, were considered as having 1 nucleus/BWM.

### Counting phenotypes in worms expressing EFF-1 in BWMs

To find whether ectopic expression of EFF-1 in BWM cells produces any special phenotypes, we used BP2137 *drh-1(tm1329)IV*; hyEx375[*myo-3p*::EFF-1, *myo-3p*::mCherry, KS bluescript]. We first isolated 4 wt-like *myo-3p::mCherry* L4 hermaphrodite worms, one worm per NGM plate+Op50. These worms were led to lay eggs and were transferred to a fresh plate 1-2 times per day during 5 days. Progeny including eggs, larvae and adults were divided into two groups, namely *myo-3p::mCherry*(+) and *myo-3p::mCherry*(-). We counted the total number of progeny with a certain phenotype (e.g. wt-like worms, adult Unc+Dpy worms, larval arrested Unc+Dpy worms and unhatched eggs) for each of the 4 mothers. Finally, we calculated the average number of worms with certain phenotype ± SEM and the fraction ± SE of worms with a certain phenotype in each group.

### Cell culture and preparation of pseudoviruses

Baby Hamster Kidney cells (BHK), BHK-21(ATCC) were cultured in Dulbecco’s Modified Eagles Medium (DMEM) and recombinant viruses were prepared as described [6] with some modifications. Briefly, BHK cells were grown to 70% confluence on 10 cm plates and then transfected using Fugene HD + OptiMEM at ratio 1:4 (Fugene:DNA), with plasmids encoding pOA20 (pCAGGS::*aff-1*::FLAG, 2µg/ml final concentration) [6] or pOA28 (pCAGGS::VSV-G Indiana, 1µg/ml final concentration)[33]. Following 24 h incubation at 37°C in 5% CO_2_, cells were infected with VSVG-complemented VSVΔG recombinant virus (VSVΔG-G) at a multiplicity of infection (MOI) of 5, for 1 hour at 37°C in a 5% CO_2_ incubator in serum free DMEM. Virus infected cells were washed 3-6 times with PBS (+ Ca^++^ & Mg^++^) to remove unabsorbed VSVΔG-G virus. Following a 24 h incubation period at 37°C, the supernatant containing the VSVΔG-G, or VSVΔG-AFF-1 pseudoviruses were harvested without scraping the cells and centrifuged at 600 g for 10 min at 4°C to clear cell debris. VSVΔG-AFF-1 Virions were filtered through 0.22 µm filter unit and then double concentrated. First, by pelleting at 100,000 g through a 20% sucrose cushion, and resuspension in 10%FBS DMEM, second, by pelleting at 100,000 g through a 10% sucrose cushion, and final resuspension in 45µl DMEM. Finally, VSVΔG-AFF-1 was incubated with anti-VSV-G antibody mAb diluted 1:1000 to inhibit infection due to residual presence of VSV-G. The effective blocking of VSV-G was confirmed by tittering pseudoviruses in BHK cells and by injection of VSVΔG-G incubated with anti-VSV-G into worms (Figure S4).

### Titering VSV pseudotyped viruses

Titering VSV pseudotyped viruses was performed as described [6] with some modifications. Briefly, 5×10^3^ BHK cells were plated into each well of a 96 well tissue culture plate (NUNC, cat# 167008). To determine the titer of VSVΔG-AFF-1, BHK cells were initially transfected with 2 µg/ml pOA20 (pCAGGS::AFF-1::FLAG). Cells transfected with empty vector served as control. Eight serial x2 dilutions of the virus were performed and added to cells. After 18-24 hours of incubation, GFP expressing cells were counted in at least three dilutions using x20 objective of Zeiss Axiovert 200M fluorescence microscope. Inoculation was performed in the presence of anti-VSV-G antibody mAb diluted 1:1000 to inhibit infection due to residual presence of VSV-G.

### Statistical tests

The specific tests used are described in the figure captions and the results section. The graphs show mean ± SEM unless noted otherwise. For each experiment at least two biological replicates were performed and the number of animals per experiment is stated in the figure legends.

## Abbreviations

AFF-1: Anchor cell Fusion Failure
BHK: Baby Hamster Kidney cells
BWM: Body Wall Muscle
Dpy: Dumpy phenotype
EC: Excretory Canal cell
EFF-1: Epithelial Fusion Failure 1
EFF-2: Epithelial Fusion Failure 2
FUSEXINS: FUSion proteins essential for sexual reproduction and Exoplasmic merger of plasma membranes
HYP: Hypodermis (epidermis)
SM: Somatointestinal Muscle
ts: temperature sensitive
UM: Uterine Muscle
Unc: Uncoordinated phenotype
VSV: Vesicular Stomatitis Virus
VSV-G: VSV-glycoprotein G
VSVΔG: pseudoviruses in which the glycoprotein G gene was deleted
VSVΔG-AFF-1: pseudovirus coated with AFF-1
VSVΔG-G: pseudovirus coated with G glycoprotein

## Acknowledgements

We thank Don Gammon for providing the *drh-1(-)* worms, Massimo Hilliard for providing *mec-4p*::mCherry, *mec-4p*::EFF-1, *odr-1p*::dsRed and *odr-1p*::EFF-1 plasmids and the Caenorhabditis Genetics Center for nematode strains. We thank Andy Fire for suggesting studying syncytial BWMs and their potential phenotypes. We also thank Sharon Inberg, Yael Iosilevskii, Rosina Giordano-Santini, Dan Cassel and Sivan Korenblit for helpful discussions and for critically reading the manuscript.

## Author contributions

A.M. and B.P. conceived the project. A.M. performed all experiments unless otherwise specified. X.L. generated *myo-3p*::EFF-1 Plasmid and BP2171-3 nematode strains and performed imaging of worms in Figure 5. E.M. improved viral preparation protocol. B.G. generated EFF-2 CRISPR plasmid and O.K. produced *eff-2(hy51)* worms. A.M. and B.P analyzed data and wrote the manuscript, with input from all authors.

## Online supplemental material

**Figure S1:**
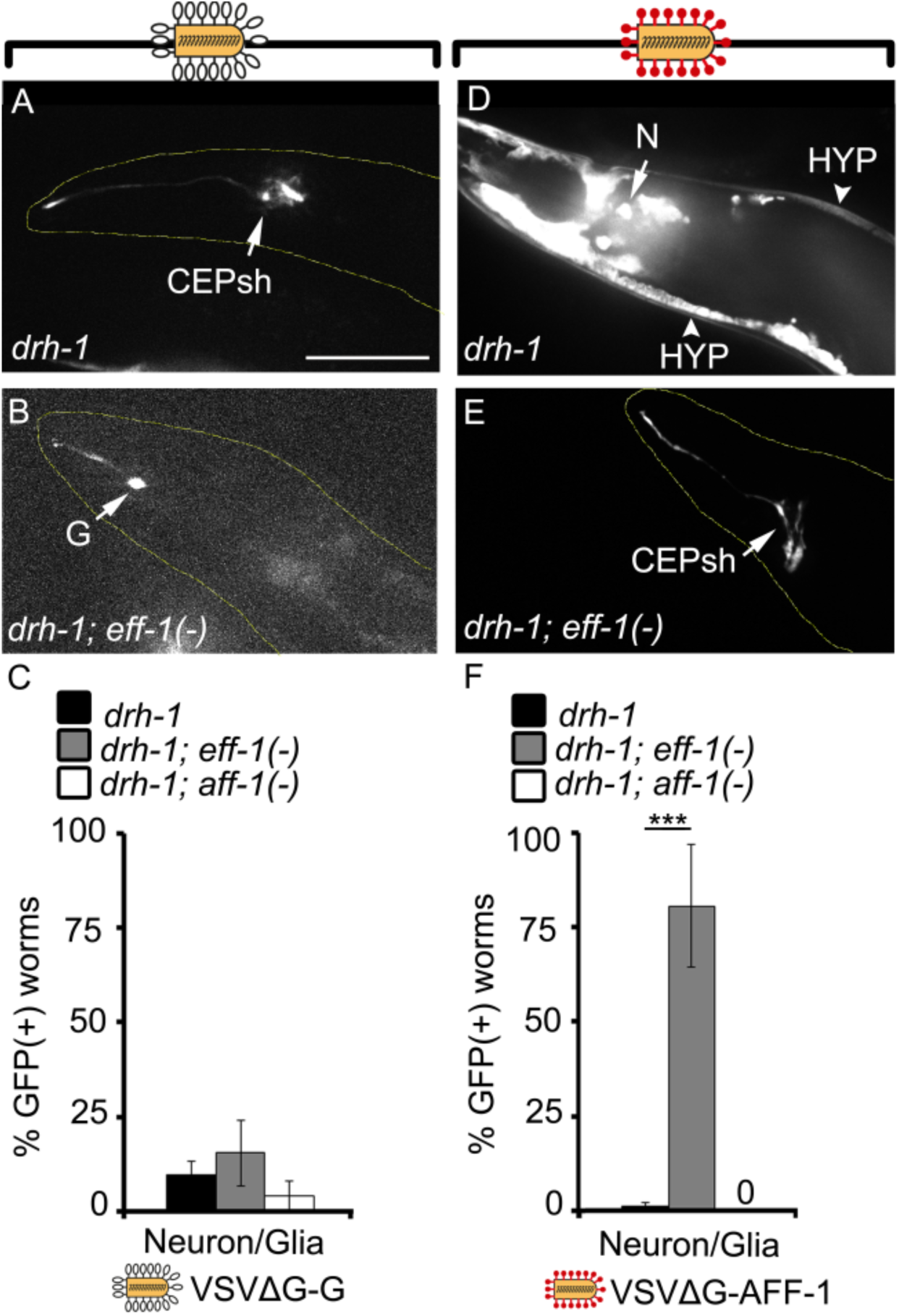
VSVΔG-AFF-1 and VSVΔG-G infect glia and neuron cells. **(A-B)** SDC microscope Z-stack projections of *drh-1* and *drh-1; eff-1(-)* worms injected in the pseudocoelom with 2300-4700 IU VSVΔG-G (white pins) and their heads imaged 48 h later. **(C)** Fraction of GFP(+) worms infected with VSVΔG-G in neuron/glia cells in the specified background. Animals were injected with VSVΔG-G and analyzed as in (A-B). **(D-E)** SDC microscope Z-stack projections of *C. elegans* cells infected with 33-240 IU VSVΔG-AFF-1 (red pins). *drh-1* or *drh-1; eff-1(-)* worms were injected in the pseudocoelom and imaged 48 h later. **(F)** Fraction of GFP(+) worms infected with VSVΔG-AFF-1 in neuron/glia cells. Animals were analyzed as in (D-E). For (A-B) and (D-E) arrowheads point to hypodermal nuclei (HYP) and arrows point to indicated infected cell. CEPsh-Cephalic Sheath (glia), G-glia, N-neuron. Scale bar, 50 µm. In (A, B and E) yellow dashed lines outline worm’s anterior body (head). In (C) and (F), bars represent average ± SEM. ***P<0.001 (Student’s T-test). For VSVΔG-AFF-1: n=46,15 and 10 for *drh-1,drh-1;eff-1(-)* and *drh-1;aff-1(-)* respectively, and for VSVΔG-G: n= 56, 39 and 15 for *drh-1, drh-1;eff-1(-)* and *drh-1;aff-1(-)* respectively).

**Figure S2:**
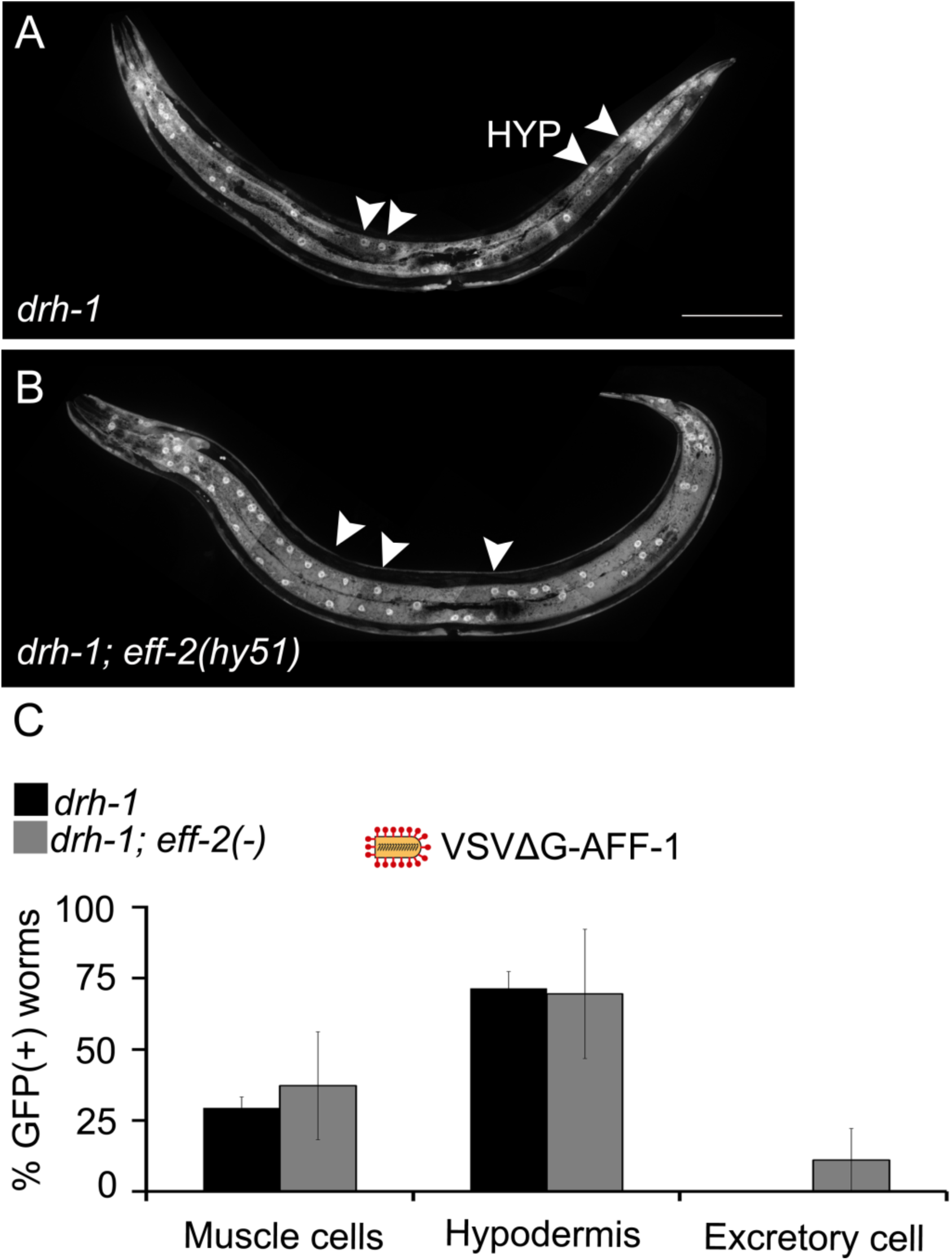
VSVΔG-AFF-1 infects *eff-2(-)* worms. **(A-B)** SDC microscope Z-stack projections of *drh-1* or *drh-1;eff-2(-)* worms injected with VSVΔG-AFF-1 (35-67 IU) and imaged 48 h later. **(C)** Fraction of GFP(+) worms infected with VSVΔG-AFF-1. Animals were injected with VSVΔG-AFF-1 and analyzed as in (A-B). Arrowheads point to hypodermal (HYP) nuclei. Scale bar, 100 µm. In (C) bars represent average ± SEM. n=23 and 20 for *drh-1* and *drh-1;eff-2(-)* respectively. For all types of infected cells, there is no significant difference between % GFP(+) worms from *drh-1* and *drh-1;eff-2 (-)* backgrounds with two-tailed Student’s t-test (p<0.05).

**Figure S3.**
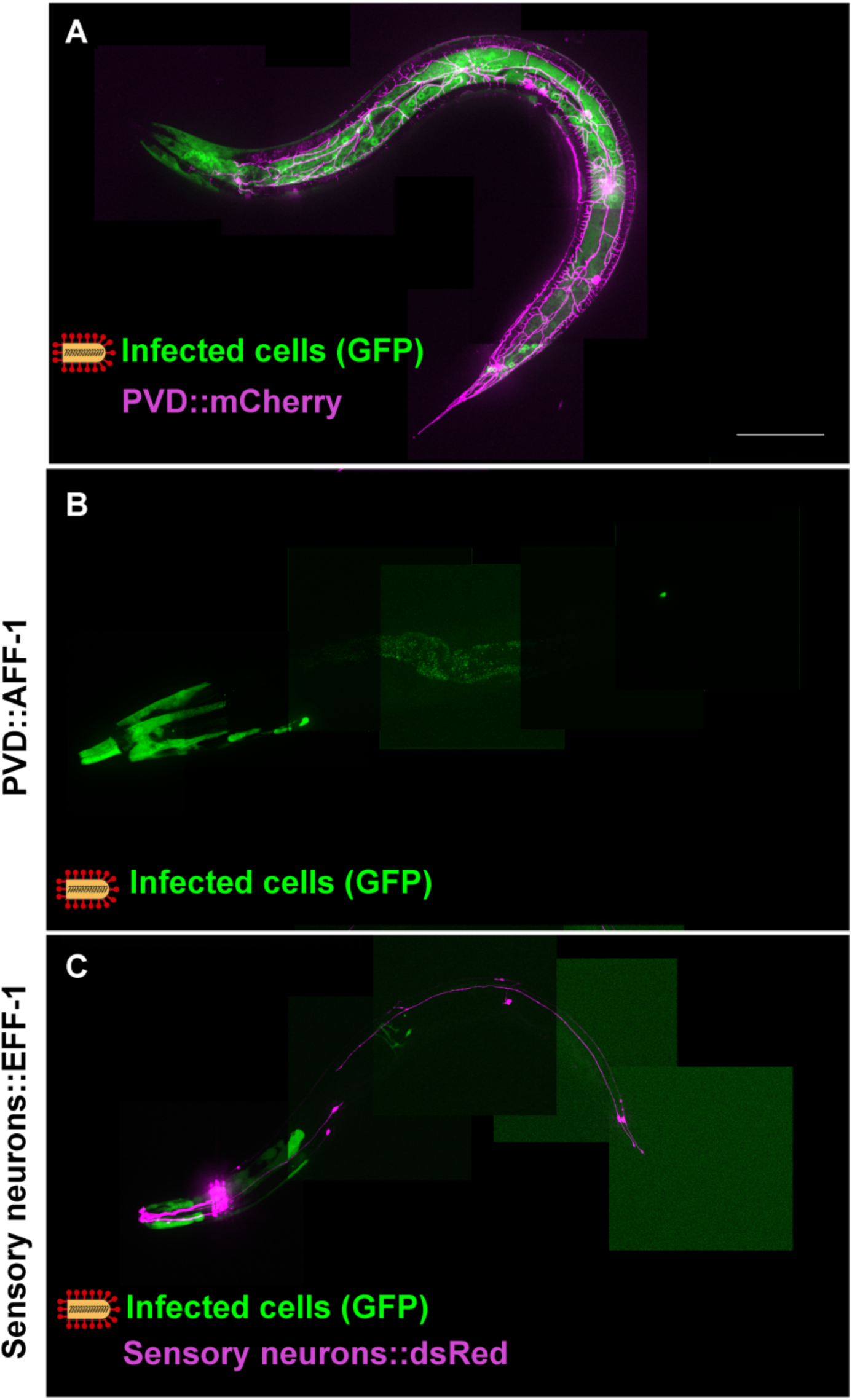
VSVΔG-AFF-1 does not infect PVD and other sensory neurons ectopically expressing AFF-1/EFF-1. **(A-C)** SDC microscope Z-stack projections of animals infected with 82-103 IU VSVΔG-AFF-1 (red pins). (For genotypes and quantitation see Table S2). Scale bar, 100 µm. **(A)** Young adult expressing mCherry in PVD. **(B)** *eff-1(ts)* adult expressing AFF-1 in PVD. **(C)** *eff-1(ts)* adult expressing EFF-1 and dsRed in 12 sensory neurons. See also Table S2 and Movie S2.

**Figure S4.**
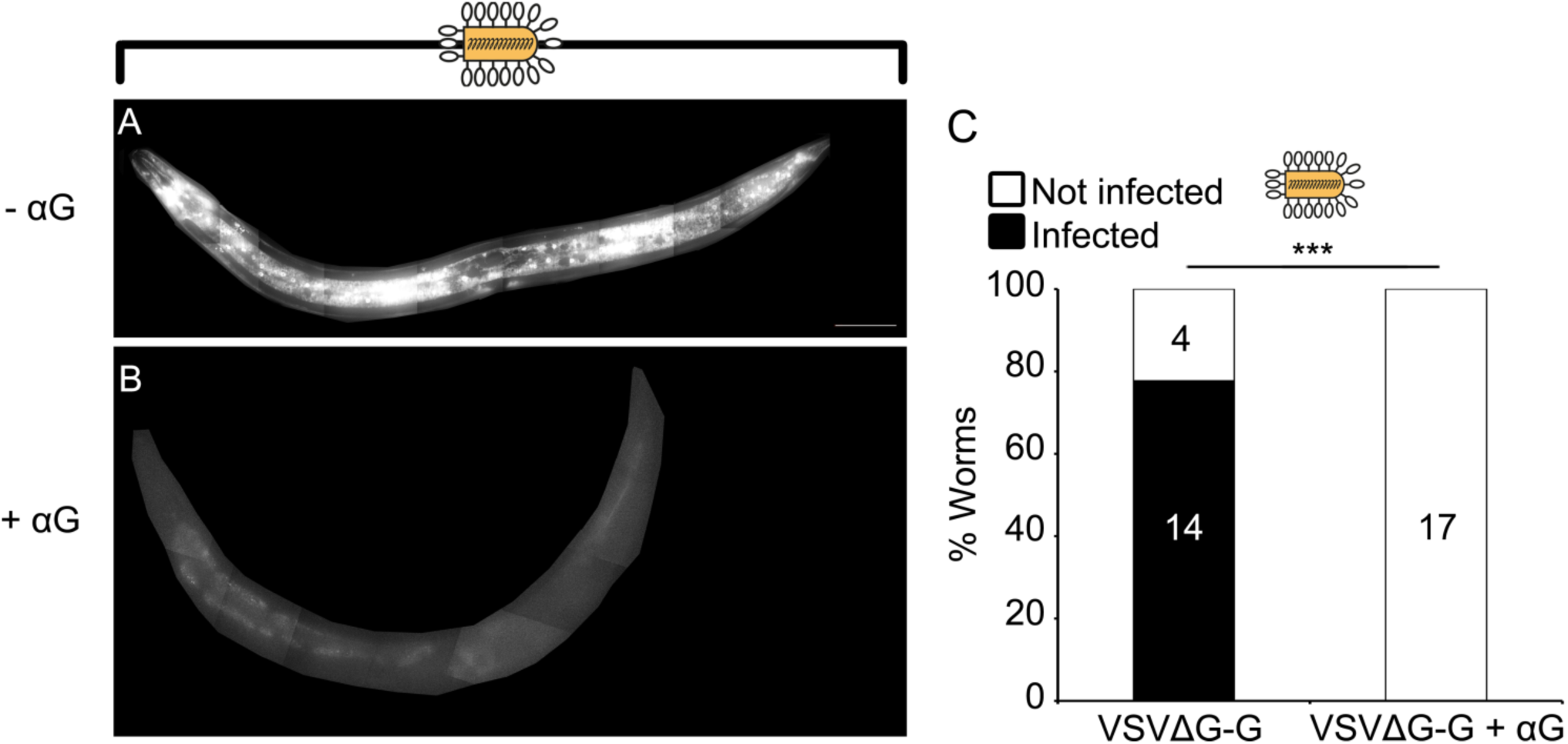
Anti-VSV-G antibody blocks VSVΔG-G infection in living worms. **(A-B)** SDC microscope Z-stack projections of *drh-1* worms injected with 4700 IU VSVΔG-G that was either not preincubated (-αG), or preincubated with αVSV-G antibody (+αG) and imaged 48 h later. Scale bar, 50 µm. **(C)** Fraction of worms that are infected or not infected with VSVΔG-G or VSVΔG-G +αG. Animals were treated as in (A-B). n=18 and 17 animals for VSVΔG-G and VSVΔG-G+αG respectively. Fisher exact test. p***<0.001.

## Supplementary tables

**Table S1.**
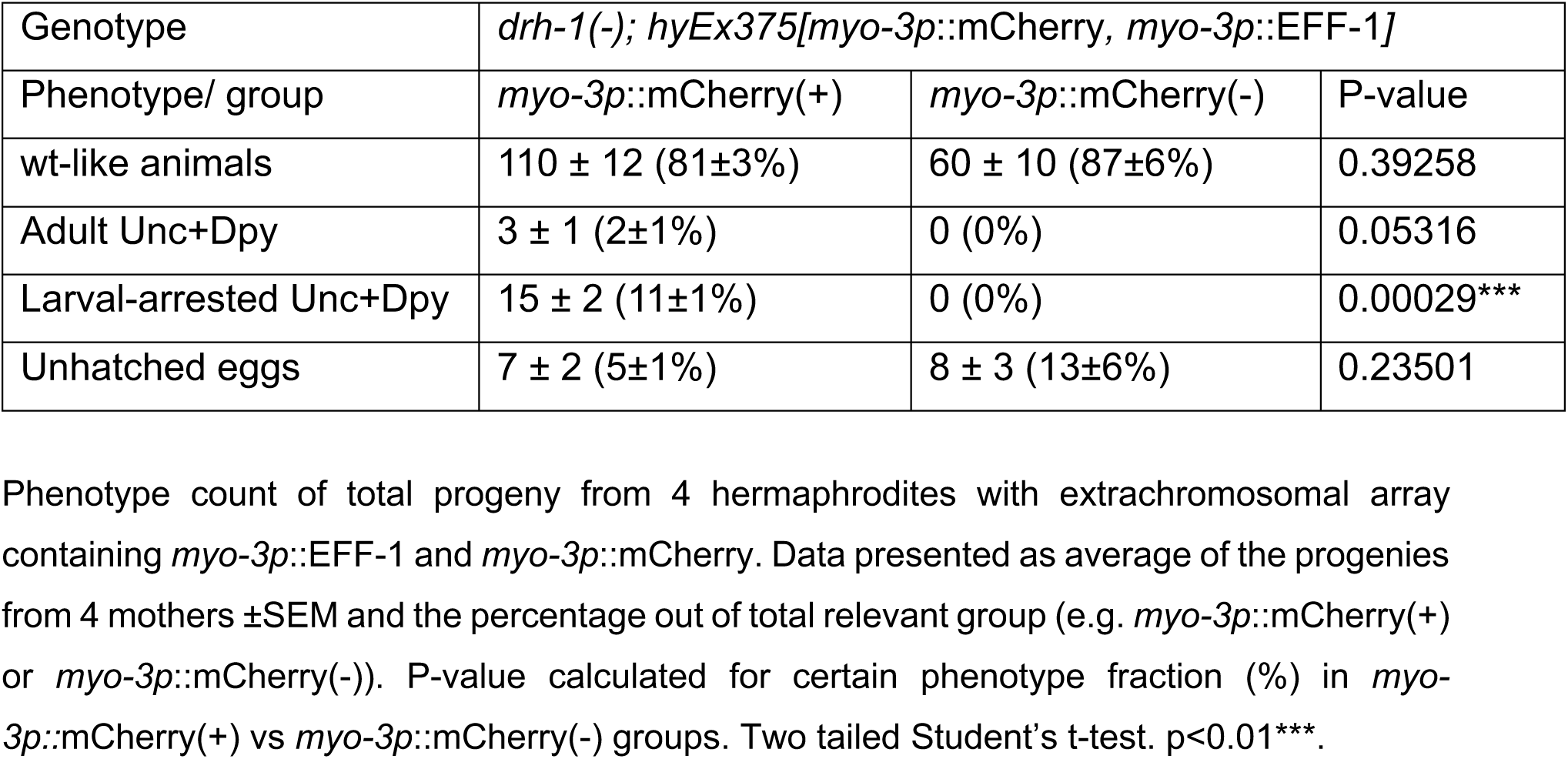
Phenotypes of worms with EFF-1 expressed in BWMs.

| Genotype | *drh-1(-); hyEx375[myo-3p::mCherry, myo-3p::EFF-1]* |  |  |
| --- | --- | --- | --- |
| Phenotype/ group | *myo-3p::mCherry(+)* | *myo-3p::mCherry(-)* | P-value |
| wt-like animals | 110 ± 12 (81±3%) | 60 ± 10 (87±6%) | 0.39258 |
| Adult Unc+Dpy | 3 ± 1 (2±1%) | 0 (0%) | 0.05316 |
| Larval-arrested Unc+Dpy | 15 ± 2 (11±1%) | 0 (0%) | 0.00029*** |
| Unhatched eggs | 7 ± 2 (5±1%) | 8 ± 3 (13±6%) | 0.23501 |

Phenotype count of total progeny from 4 hermaphrodites with extrachromosomal array containing *myo-3p::EFF-1* and *myo-3p::mCherry*. Data presented as average of the progenies from 4 mothers ±SEM and the percentage out of total relevant group (e.g. *myo-3p::mCherry(+)* or *myo-3p::mCherry(-)*). P-value calculated for certain phenotype fraction (%) in *myo-3p::mCherry(+)* vs *myo-3p::mCherry(-)* groups. Two tailed Student's t-test. p<0.01\*\*\*.

**Table S2.** AFF-1/EFF-1 expression in sensory neurons does not induce their infection with VSVΔG-AFF-1.

| Strain | Genotype | Description | total # of worms | # VSVΔG-AFF-1-infected worms | # worms with GFP(+) cells overlapping with EFF-1/AFF-1 expressing cells |
| --- | --- | --- | --- | --- | --- |
| BP2126 | *drh-1(tm1329); eff-1(hy21); hyEx373 [pmyo-2::GFP, pdes-2::AFF-1(pME4), KS]* | AFF-1 expressed in PVD (Oren-Suissa et al 2017) , in *eff-1(ts)* background | 18 | 6 | 0 |
| BP2131 | *drh-1(tm1329); eff-1(hy21); hyEx374 [pmec-4::EFF-1, pmec-4::dsRed, podr-1::EFF-1, podr-1::dsRed, KS] line1* | EFF-1 expressed in 12 sensory neurons (kind gift from M. Hilliard lab), in *eff-1(ts)* background | 28 | 15 | 0 |
| BP2132 | *drh-1(tm1329); eff-1(hy21); hyEx374 [pmec-4::EFF-1, pmec-4::dsRed, podr-1::EFF-1, podr-1::dsRed, KS] line2* | EFF-1 expressed in 12 sensory neurons, in *eff-1(ts)* background | 22 | 7 | 0 |
| BP2133 | *drh-1(tm1329); eff-1(hy21); hyEx374 [pmec-4::EFF-1, pmec-4::dsRed, podr-1::EFF-1, podr-1::dsRed, KS] line3* | EFF-1 expressed in 12 sensory neurons, in *eff-1(ts)* background | 35 | 13 | 0 |

**Table S3.**
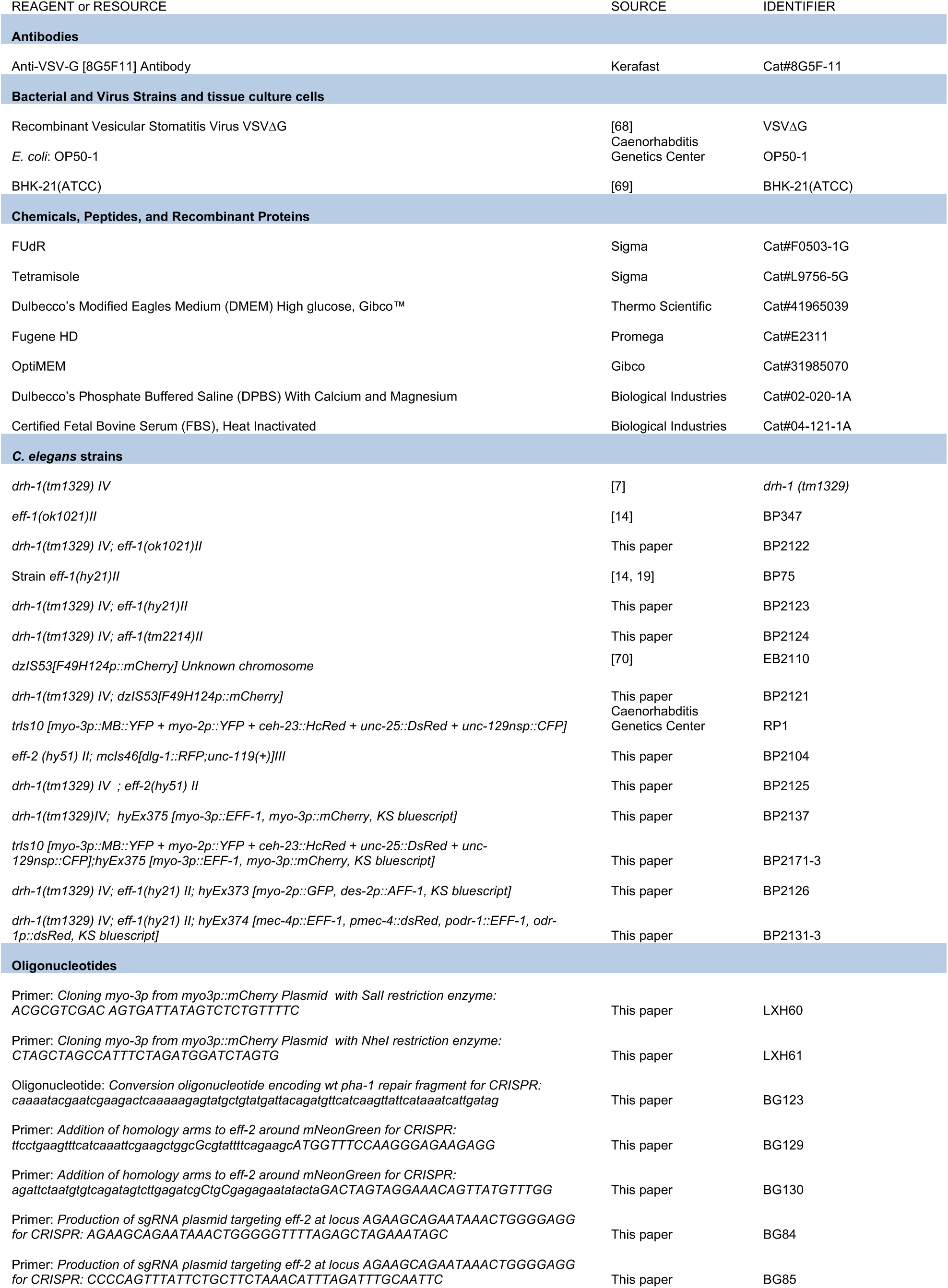
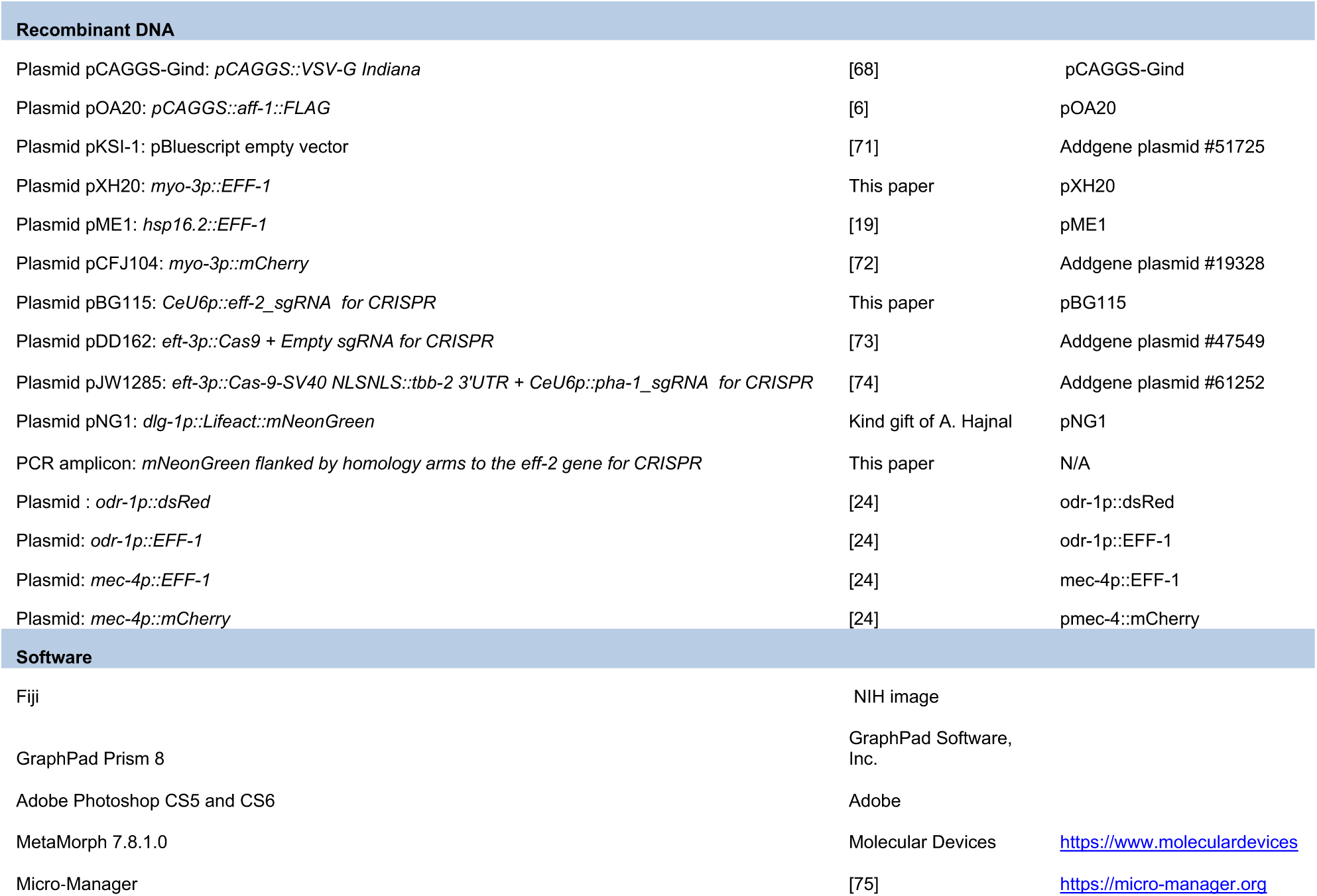
Reagents, strains, oligos, plasmids and software.

## Data for movies

**Movie S1.**
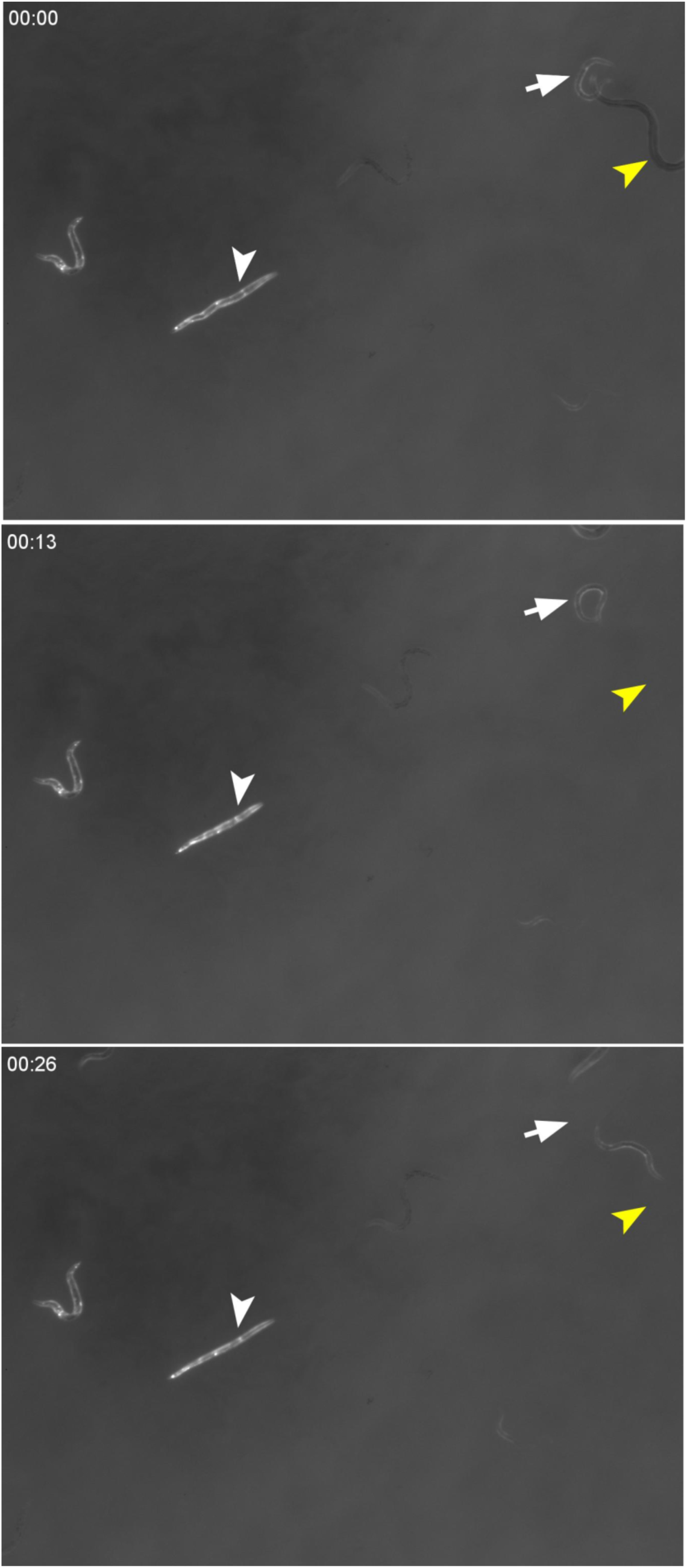
EFF-1 ectopic expression in BWMs results in Uncoordinated and Dumpy (Unc+Dpy) phenotypes. Mixed population of worms with extrachromosomal *pmyo-3::mCherry* and *pmyo-3::EFF-1*. White arrowhead point to mCherry(+) worm that is Unc+Dpy. White arrow point to mCherry(+) worm that is wt-like. Yellow arrowhead points to a wt-like mCherry(-) worm, which left the frame within seconds. Elapsed time (seconds) indicated in top left corner.

**Movie S2.**
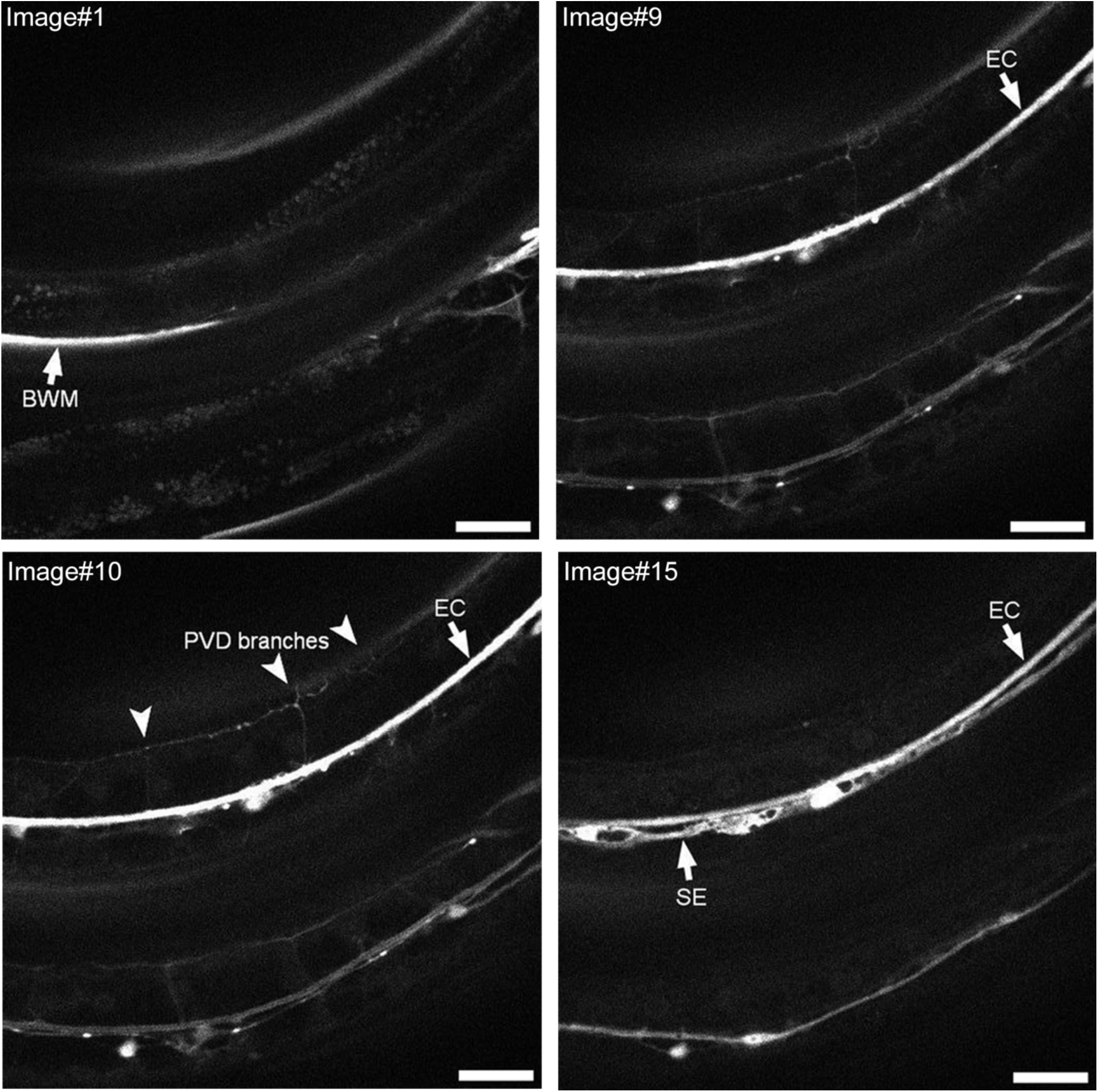
VSVΔG-G infects PVD neuron. SDC microscope Z-stack of young adult worm injected with 10^6^ IU VSVΔG-G and imaged 48h later. Arrowheads, infected PVD’s candelabra/menorahs arborized branches. Arrows, infected cell. BWM-Body Wall Muscle, EC-Excretory Cell, SE-Seam cell syncytium. Scale bars, 25 µm. Note that there is a second worm (bottom, not indicated in images), also showing infected PVD branches, EC, SE and a muscle cell.

